# Rarefaction is better than robust Aitchison PCA and other compositional data analysis methods at controlling for uneven sequencing effort

**DOI:** 10.64898/2026.01.06.697977

**Authors:** Patrick D. Schloss

## Abstract

Amplicon sequencing typically results in a wide distribution in the number of sequences obtained from each sample. How best to account for this variation has been a persistent problem in the microbial ecology literature. Historically, rarefaction has been used in ecology and this practice was adopted by microbial ecologists. But rarefaction has been strongly criticized leading to the development of compositional data analysis and normalization methods. Therefore, I reassessed the benchmarking data that was generated by the developers of one such method, robust Aitchison PCA. I found numerous problems in the Python code that led to the support of robust Aitchison PCA. These problems extended to the creation of simulated datasets, implementation of machine learning methods, and choice of analysis parameters. Furthermore, the analysis of the simulated and case study datasets was done in a manner that was foreign to standard microbiome analyses. I corrected the problems in the original code, added datasets with smaller effect sizes, and expanded the collection of methods that purport to correct for uneven sequencing effort. Contrary to the claims of the original analysis, robust Aitchison PCA was not insensitive to uneven sequencing effort and did not perform as well as rarefaction. In fact, even using the benchmarking framework from the original analysis, rarefaction outperformed robust Aitchison PCA, other compositional data analysis methods, and other normalization methods. Rarefaction remains the preferred method of controlling for uneven sampling effort in amplicon sequence studies.

**Importance:** Efforts to connect the structure of microbial communities with environmental processes and host health have captured widespread interest among scientists and the general public. The methods used to generate the sequencing data that are fundamental to these efforts result in wide variation in the number of sequences per sample. This variation and how to account for it can have detrimental effects on the ability to draw valid conclusions. Compositional data analysis methods including robust Aitchison PCA have grown in popularity for mitigating these effects and have been proposed as alternatives to rarefaction. In this study, I reviewed and fixed the code that was used to benchmark robust Aitchison PCA and expanded the analysis. With this improved and expanded analysis, I found that rarefaction was still superior to robust Aitchison PCA and other methods of controlling for uneven sequencing effort.

## Introduction

A persistent challenge in amplicon and shotgun metagenomic sequencing of microbial communities is the variation in the number of reads obtained across samples (e.g., 1, 2, 3). The variation in the number of reads generally occurs because of an accumulation of sampling errors during attempts to pool equimolar amounts of DNA to generate libraries for subsequent sequencing. It is not uncommon to observe as much as 100-fold variation in sequencing depth (4). This challenge is not unique to microbial ecologists and has long plagued landscape and animal ecologists (5, 6). The classical approach to control for uneven sampling has been rarefaction (7). In rarefaction, a consistent number of individuals are randomly sampled from each sample. An alpha or beta diversity metric is then calculated from the sample. The random subsampling and diversity measurement steps are repeated a large number of times (e.g., 100 or 1,000) and the measurements are averaged over the subsamples (4). Unfortunately, the term “rarefying” and occasionally rarefaction have been misidentified as a single subsampling step (4, 8). Regardless, over the past 10 years, the process of rarefaction has become controversial because some view the choice of the sampling depth as arbitrary, the metrics can become more similar than they truly are, and because the practice of downsampling to a common number of sequences leaves the impression that good data are being unnecessarily discarded (8, 9). Numerous alternatives have been proposed including calculating the relative abundance of each taxon in a community so that they sum to 1 within a sample (10), using a single subsampling step (11), normalization of counts so that the total number of individuals sums to a user defined value (12, 13), transforming the data using methods commonly used for gene expression analysis (8, 14), use of estimators of diversity that can extrapolate beyond the number of observed individuals (15–17), and the use of compositional statistics based on a log ratio of taxon counts to a reference value (10, 18–23).

Researchers have adopted compositional methods because the number of sequences obtained per sample is typically independent of the true number of individuals in a community and so the number of times each taxon is observed is not independent of the other taxa (10, 23). The general approach is to calculate a centralized log ratio (i.e., CLR) where the abundance of each taxon within a sample is divided by the geometric mean abundance within the sample, which is then used as the argument for the natural logarithm. These values are then used to calculate pairwise Euclidean distances between samples resulting in so-called Aitchison distances (24). Aside from controlling for the compositional nature of sequence data, proponents of the Aitchison distance claim that the distances are insensitive to differences in sampling effort (10, 23). A challenge of Aitchison distances is that because microbial communities are patchy a large number of features often have a frequency of zero. These zero values cause the geometric mean to be undefined. To overcome this challenge, factors are added to the data. Three approaches have been proposed: adding a pseudocount of 1 all frequency data (i.e., One CLR), adding a pseudocount of 1 divided by the total number of individuals in the sample (i.e., Nudge CLR), and imputing zero values to be a random small value (i.e. Zero CLR) (20–22). As an alternative to these transformations, Martino et al. (23) (i.e., Martino) proposed calculating CLR values for each sample after ignoring zero values followed by matrix completion using the OptSpace matrix completion algorithm. They called this method robust Aitchison principal components analysis (PCA) (i.e., RPCA). The authors claimed that the advantage of RPCA over the other CLR-based approaches was that it did not require imputing zero values and that the method was less sensitive to uneven sampling effort.

In contrast to these claims, I have shown that the output of RPCA and other alternatives to rarefaction are more sensitive to uneven sampling than rarefaction (4, 25). Furthermore, use of these methods can inflate the false detection rate and decrease the statistical power to detect differences in alpha and beta diversity metrics. Given the near unanimous agreement over the past decade that rarefaction is “inadmissible” (8) and the development of alternatives to rarefaction (12–14, 20–23), one might question why my results disagree so markedly with those of others. There are several possible reasons. First, researchers often propose new methods by demonstrating their method reveals new insights over old methods without objectively benchmarking the method. Because one cannot know the true difference between samples, a positive or negative result with experimental data cannot be interpreted. Second, when benchmarking tests are developed, researchers may inadvertently select tests that favor the method they are developing. This proved to be the case for the original paper that called rarefaction into question (8, 25). Third, the choice of how to simulate community data can inadvertently favor the method being developed. A popular aphorism states that “all models are wrong, some are useful” (26). Similar to one’s choice of benchmarking tests, the modelling approach may favor some methods over others. For instance, I have critiqued how others have accounted for variation in sampling depth as being unlike what we see in experimental data (25). To mitigate against bias, it is necessary to use a comprehensive benchmarking test suite that allows one to compare diverse methods using inference methods that are commonly used by researchers.

In my previous analyses advocating for rarefaction, I generated null community models and treatment groups with known effect sizes by creating taxon empirically-determined abundance distributions and sampling depths based on 12 published datasets (4). Alternatively, Martino modelled communities with known effect sizes using a statistical model that sought to mimic the diversity of communities described in a study comparing the microbial communities found on people’s fingertips and their computer keyboards (27). As described in Martino, the overall simulation framework involved simulating two treatment groups each represented by 100 communities for a total of 200 communities. Each community contained as many as 1,000 taxa. They stated that for each treatment group they fit a normal distribution to the relative abundances of taxa from two individuals’ samples to obtain a mean and standard deviation for each taxon. The fraction of taxa that overlapped between treatment groups was also selected to mimic the overlap observed between the two individuals. The expected taxon relative abundances for each sample were the same across each sample within the same treatment group. Next, a common normal distribution with a single mean and standard deviation was used to generate noise (i.e., homoscedastic noise) that was added to each relative abundance value to create variation across samples and features. Next, a random subset of sample-taxa combinations were selected for incorporation of additional noise (i.e., heteroscedastic noise). Finally, the sample-taxa combinations were randomly sampled using a Poisson-lognormal distribution to generate the simulated number of sequences for each sample-taxa combination so that the total number of sequences within a sample contained approximately 1,000, 2,000, 4,000, or 10,000 sequences per simulation; one set of 200 samples were simulated for each sequencing depth. In addition to the simulation where a subset of taxa was shared in both treatment groups, Martino created positive and negative control simulations. In the positive control simulation, there were no overlapping taxa resulting in two clearly defined groups of samples. In the negative control simulation all the taxa were shared in similar abundance so that the groups of samples could not be distinguished. Finally, they compared RPCA to other methods using two published datasets.

This difference in modelling strategy created an interesting opportunity to reassess the performance of RPCA and other CLR-based approaches relative to rarefaction. Although Martino used an interesting modelling approach, they did not compare RPCA to other CLR-based approaches or to rarefaction. Furthermore, they used benchmarking tests that did not reflect how researchers typically analyze microbial community data. The goal of the current study was to repeat the benchmarking of RPCA and other CLR-based methods relative to rarefaction using the modelling framework described by Martino.

## Results

### Description of simulated communities

To maintain a connection to Martino’s original simulations and implement a broader set of tests, several changes were necessary in the implementation of Martino’s code. First, Martino’s code did not return the same number of sequences for each sample. Because one goal of the current study was to assess the effect of variation in sequencing effort on the performance of RPCA, I modified the parameters inputted to Martino’s code to produce more than the desired number of sequences per sample and then performed a single subsampling of the samples to 1,000, 2,000, 4,000, or 10,000 sequences per sample to have exactly the desired number of sequences per sample. Second, to account for the effect of the random number generator, instead of replicating each simulation condition only once as Martino did, I replicated each condition 10 or 100 times depending on the analysis. Third, Martino only analyzed their positive and negative control simulations with 4,000 sequences. My implementation of their simulations and subsequent tests added the sequencing depths of 1,000, 2,000, and 10,000 sequences for the positive and negative controls and for the simulations with partially overlapping treatment groups. The regenerated heatmaps and PCoA plots in Figure 1 are qualitatively similar to those in Figure 3 of Martino suggesting that my changes to the original code were superficial. The most notable difference to Martino’s version of the figure was that they inadvertently switched the labels of the axes and performed the One CLR transformation on features rather than samples (Supplementary Text). Replicating what Martino reported, the first two axes of the Bray-Curtis ordinations only explained a fraction of the total variation in the data while the RPCA ordinations explained all the variation. However, this was because RPCA is run by giving the algorithm a desired number of axes; in this case, 2 axes. Regardless, the ratio of the fraction of the variation explained by the first two axes were similar between RPCA and PCoA with Bray-Curtis.

**Figure 1.**
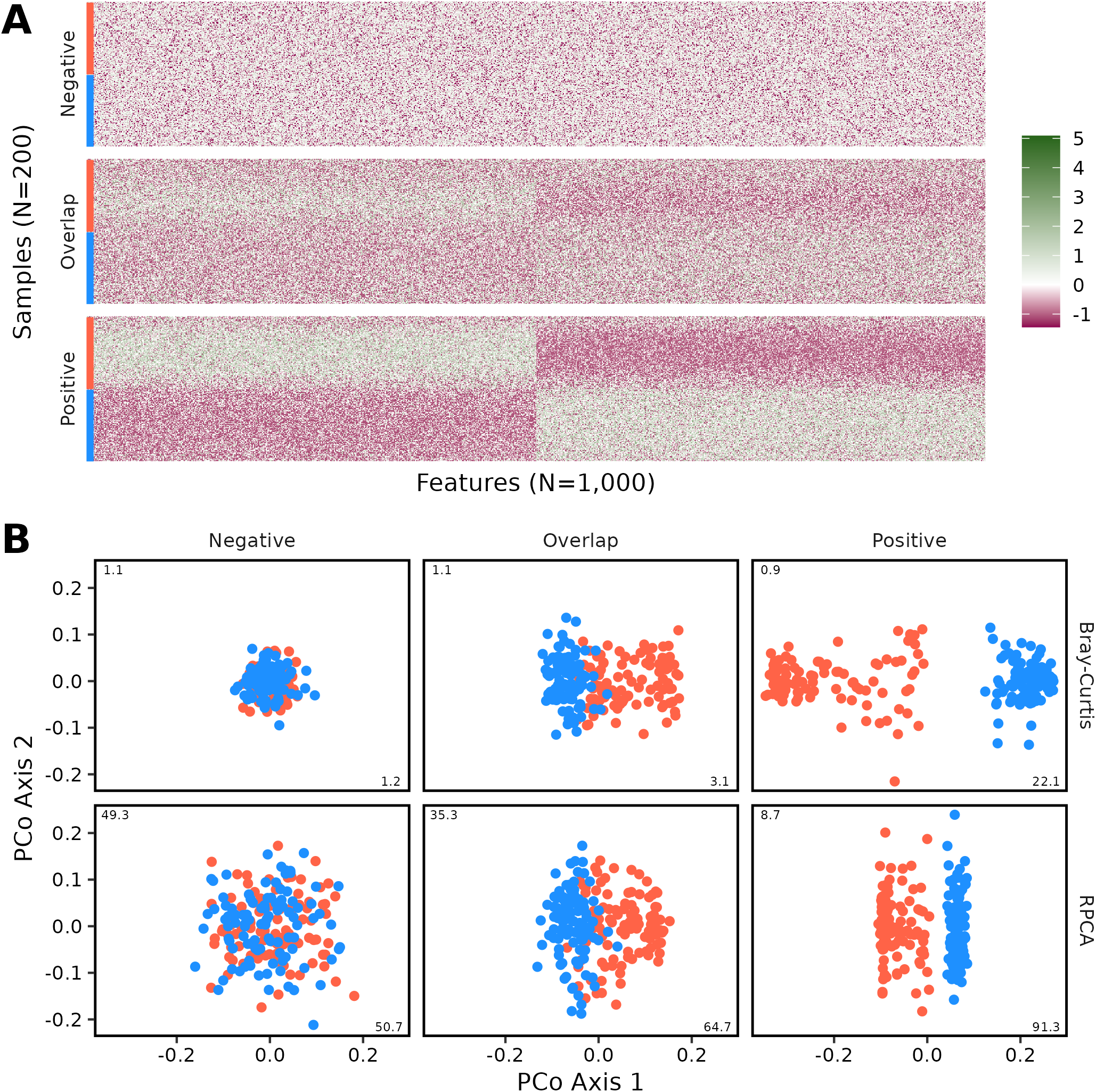
Rerunning the original Martino code in my computational environment produced qualitatively similar heatmaps (A) and ordination diagrams (B) to what they originally reported when 4,000 sequences were sampled from each sample. This figure is analogous to Figure 3DE of Martino with the inclusion of additional data from simulations where there was partial overlap between the two treatment groups (“Overlap”). The red and blue bars on the left edge of the heatmaps in panel A indicate the treatment group that each sample belonged to. The heatmaps display the One CLR transformation of sequence counts. The PCoA diagrams in panel B were generated using Bray-Curtis distances. The numbers on the axes of each ordination diagram indicate the percent of the total variation that each axis explained.

### Comparison of simulated communities using PERMANOVA

Martino used the PERMANOVA F statistic to compare the ability of RPCA to differentiate treatment groups using raw counts or removing zero values prior to calculating the CLR (RCLR). Using the partially overlapping treatment groups they found that the RCLR pretreatment yielded larger F statistics than using raw counts. Repeating Martino’s PERMANOVA analysis, I was able to replicate their results when using RPCA (Figure 2). However, by generating 10 replicate datasets (Martino only performed one), I observed considerable variation across replicates for the F statistics calculated using RPCA with raw counts. Unfortunately, Martino did not make any comparisons to other methods of calculating distances. When I calculated Euclidean distances using other CLR and DeSeq2 transformations and Bray-Curtis distances using raw counts and normalized data, all resulted in small F statistics. I suspected that the RPCA data generated relatively large F statistics because the Euclidean distances calculated using RPCA was only done with the first two axes. This would have removed a considerable amount of variation that was embedded in the third and higher axes. The result would be elevated F statistics. To test this hypothesis, I performed PCoA on Bray-Curtis distances, calculated Euclidean distances using the first two axes of the resulting PCoA, and subjected those distances to PERMANOVA (see “Bray-Curtis PCoA” data in the Euclidean distance section of Figure 2). The calculated F statistics were even larger than those generated by RPCA. This suggested that RPCA may make communities appear more similar than they are when differences are more pronounced in higher axes. Regardless of the relative scale of the F statistics, the replicates for all approaches yielded significant P-values except for those generated using RPCA with raw counts.

**Figure 2.**
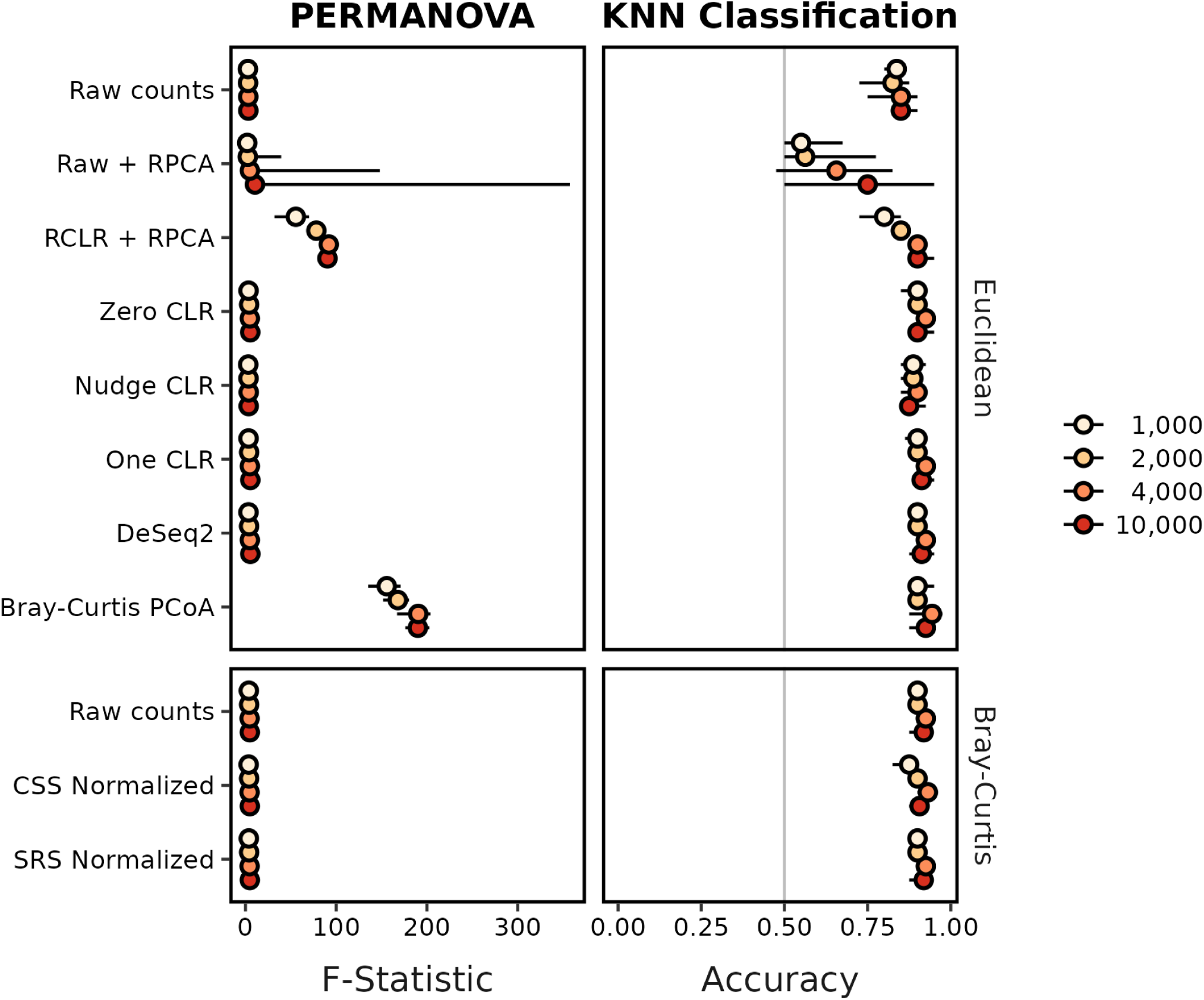
RPCA-based methods did not consistently perform as well as other methods by PERMANOVA or KNN classification when attempting to distinguish between overlapping communities. This figure includes data that was shown in in Figure 3BC of Martino with the inclusion of results when using additional data transformations, Bray-Curtis distances, and I used 10 instead of 1 replicate simulation. Each circle represents the median of ten replicate simulations and the confidence line indicates the range between the minimum and maximum values. The median KNN classification accuracy and range were the median of 10 median values each calculated across 100 training-testing splits using the fitted K value from 80:20 splits. The vertical gray line for KNN classification indicates the accuracy one would expect from randomly guessing the treatment group.

### Comparison of simulated communities using K-nearest neighbors (KNN) Classification

In addition to comparing PERMANOVA F statistics, Martino used KNN classification to distinguish between the two partially overlapping treatment groups. Although Martino’s text indicated they used 40% of the samples in the training set and 60% in the testing set (i.e., a 40:60 cross validation split), their code indicated that the design was actually the opposite with 60% of the samples in the training set and the remaining in the testing set (Supplemental Text); a 60:40 split is more conventional. To execute the KNN classification, Martino used the first two axes calculated from RPCA using raw counts and RCLR transformed data as the input to their KNN classification. I replicated the 60:40 split 100 times for each of the 10 simulations reporting the median accuracy on the 40% testing data for each simulation. The original Martino data indicated that the accuracy of using raw counts with RPCA was near 0.5, indicating it did no better than randomly guessing. While the minimum values I observed across the 10 replicate simulations overlapped with these results, the median values were all better than random (“Raw + RPCA” in Figure 2). I was able to replicate Martino’s result showing that using RCLR transformed counts as input to RPCA yielded higher accuracy than using raw counts.

Next I sought to place the RPCA results in context with other methods (Figure 2). I generated Euclidean distance matrices using other CLR methods and DeSeq2. I also generated Bray-Curtis distance matrices using the raw counts and CSS and SRS Normalization. These methods all yielded similar results, likely because all of samples in the simulations had the same sequencing depth. Compared to these other approaches, RPCA varied the most by sequencing depth. Although accuracy values across all methods were comparable at 4,000 and 10,000 sequences, the RPCA-based methods underperformed the others at 1,000 and 2,000 sequences.

Martino did not report the classification accuracy for their positive and negative controls. As would be expected, the mean accuracy for classifying the treatment groups in the negative control model across each depth, transformation, and distance calculation was 0.50 (sd = 0.01). Furthermore, the mean accuracy for classifying samples in the positive control was 0.99 (sd = 0.01).

Rather than using a training data set to identify an optimal number of neighbors (i.e., K), Martino used 13 neighbors. To assess whether the classification was sensitive to the number of neighbors, I split the 60% of samples in the training data into another 60:40 split to train and optimize the number of neighbors 100 times (28). Across the various conditions, the median optimal number of neighbors was 33 (IQR=21 to 41). When I repeated the search for the optimal value of K using a more conventional 80:20 split, the median optimal number of neighbors was 58 (IQR=42 to 65). Regardless of finding that the optimal number of neighbors was consistently larger than the 13 that Martino used, the difference in accuracy values was only modestly better than the other fitting procedures when using 80:20 splits with the simulated data (Figure S1).

### Comparing PERMANOVA and KNN classification results with more similar treatment groups

Using the simulation where the treatment groups partially overlapped, the PERMANOVA analyses yielded significant P-values and KNN classification were effective at distinguishing between the groups for nearly all of the methods. Because the simulated effect size was so large, it was not possible to easily differentiate between the methods’ performance. Therefore, I created treatment groups that were more similar than those simulated by Martino. Unfortunately, in the process of modifying their code to decrease the effect size, I identified a number of incongruities between Martino’s written description in their paper and in their code. This limited my ability to reuse their code to simulate greater overlap between the treatment groups (Supplemental Text). These issues included: (i) samples rather than features were shared between treatment groups; (ii) among samples from the same treatment group, the relative abundance distributions for the features were uniform; (iii) the total number of individuals in the samples varied across the simulated datasets; (iv) the heteroscedastic source of noise was actually applied to all sample-feature combinations; (v) negative mean values were used when sampling normal distribution sampling functions when calculating the heteroscedastic noise; (vi) negative abundance values generated after applying the heteroscedastic noise were made positive; and (vii) setting the random number generator seed was unable to produce reproducible simulations. I rewrote the simulation code in the R programming language to correct these issues. To simulate a more nuanced difference in treatment groups, I increased the total number of features across all samples from 1,000 to 10,000 and set 12 features to be differentially abundant between the treatment groups. These parameter values were selected to create conditions where approximately 50% of PERMANOVA tests using Bray-Curtis distances yielded P-values less than 0.05. Using the same number of features, I recreated negative and positive control simulations in which all features were shared or where the features were differentially represented between one of the two treatment groups. The resulting simulations provided benchmarking data that is more indicative of what is commonly observed with empirical data than what was observed by Martino (Figure 3). The simulations were replicated 100 times to obtain greater precision in calculating the power to detect differences by PERMANOVA and the accuracy of KNN classification.

**Figure 3.**
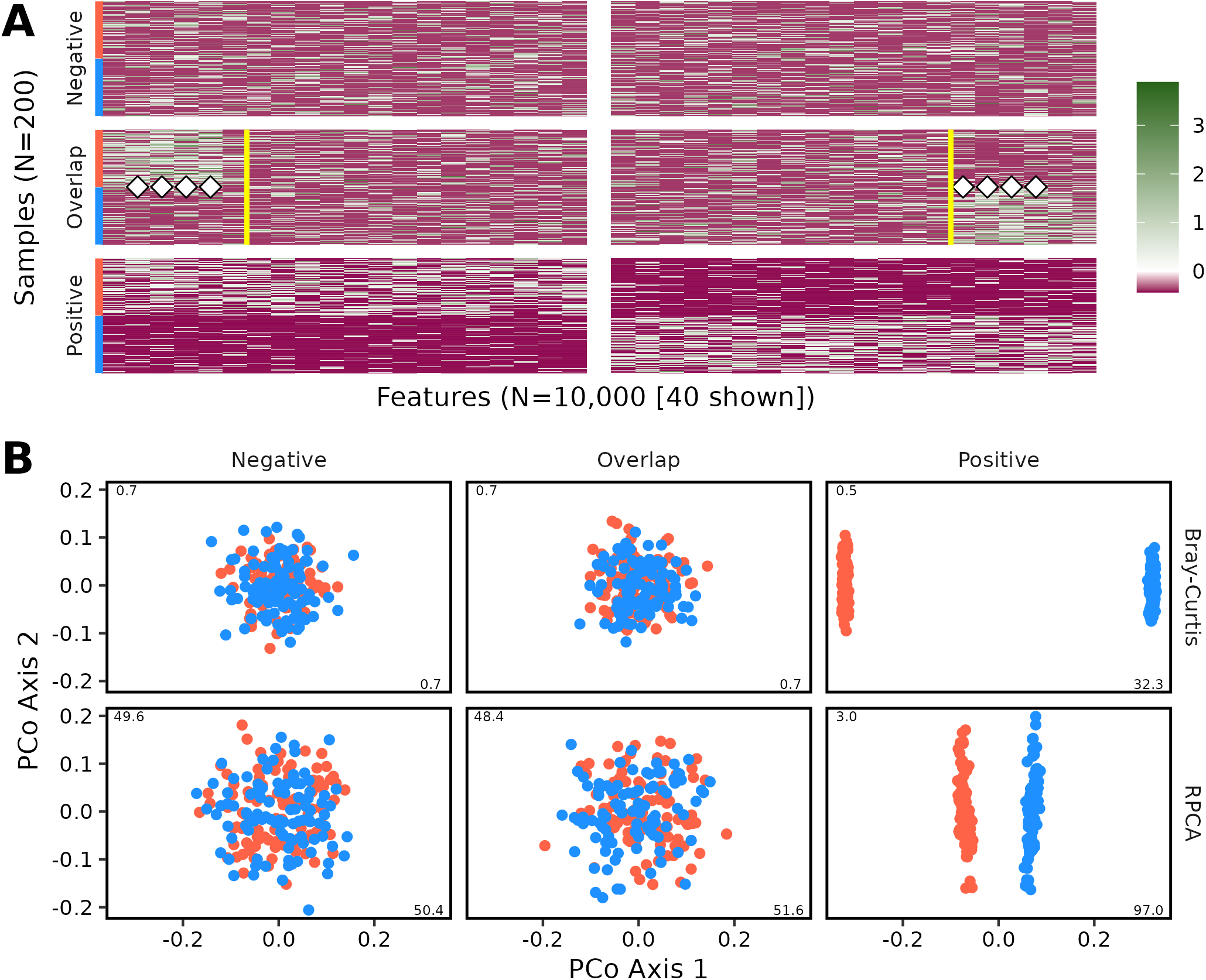
Heatmaps (A) and ordinations (B) of an example dataset comparing communities simulated rewritten in R based on the description in Martino when 10,000 sequences were sampled from each sample. This figure is analogous to Figure 1. To highlight differences in overlap simulation, only the first and last 20 features from each sample are shown in the heatmaps of panel A. The vertical white band indicates the break in the data presentation. The yellow vertical lines on the heatmap indicate the 12 features that were differentially represented in the two treatment groups. The white diamonds indicate features that had significantly different relative abundances when tested by a Wilcoxon test with P-values corrected using a Benjamini-Hochberg correction. For this example, the differences between the positive control (P=0.001) and overlap dataset (P=0.018) were significant by PERMANOVA and non-significant for the negative control (P=0.50) using Bray-Curtis distances and significant for the positive control (P=0.001) and non-significant for the overlap dataset (P=0.11) and negative (P=0.49) control using RPCA.

Using the revised simulation data revealed a considerable difference in the performance of PERMANOVA and KNN classification relative to the original simulation. Among the various methods, the RPCA-based methods performed the worst by both PERMANOVA and KNN classification (Figure 4). There was effectively no statistical power to detect differences using RPCA or to classify samples using KNN classification even at the largest sequencing depths. It was not possible to distinguish the performance between the various CLR and DeSeq2 methods using Euclidean distances or any of the methods when Bray-Curtis distances were calculated. Among these methods, the power to detect differences was routinely above 0.75 when 10,000 sequences were generated per sample. For these methods, there was a clear relationship between the number of sequences obtained from each sample and the observed power and accuracy. Similar to the earlier comparison of approaches for fitting the number of neighbors, the performance of KNN classification was modestly better using 80:20 splits with a grid-based search (Figure S2). From the revised simulations, it is evident that the performance of RPCA was considerably worse than the other methods.

**Figure 4.**
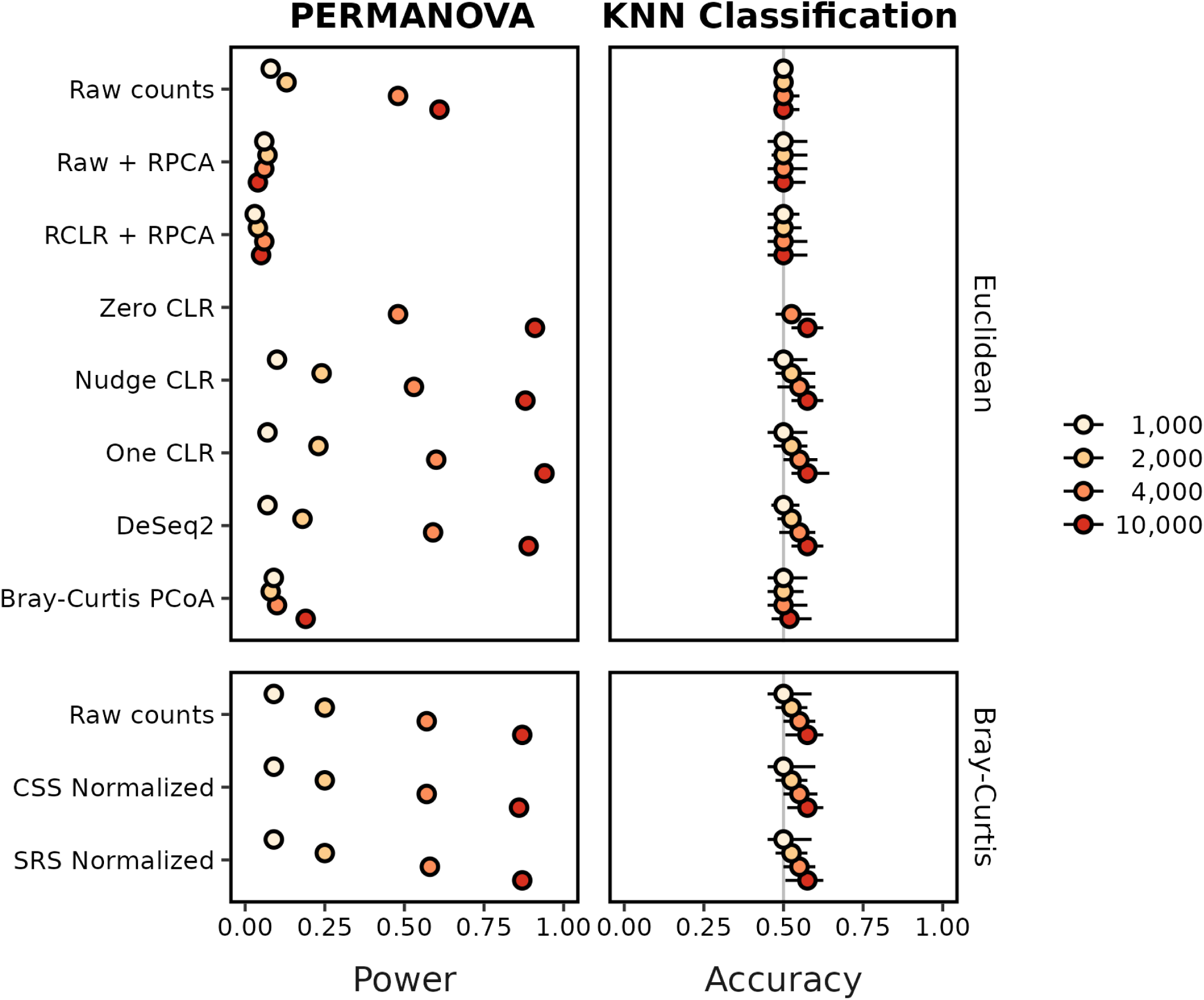
Relative to other methods, RPCA-based methods were not as effective at differentiating between groups of communities using PERMANOVA and KNN classification when using a revised simulation approach. This figure is analogous to Figure 2 except that it uses 100 instead of 10 replicates and the power to detect differences by PERMANOVA is shown instead of the F statistic. Each circle represents the median of the 100 replicate simulations and the confidence line indicates the range between the minimum and maximum values. The median KNN classification accuracy and range were the median of 10 median values each calculated across 100 training-testing splits using the fitted K value from 80:20 splits. The vertical gray line for KNN classification indicates the accuracy one would expect from randomly guessing the treatment group. The power and accuracy values for 1,000 and 2,000 sequences per sample when using Zero CLR were not calculated since the data were too sparse for the imputation algorithm to complete.

### Description of case study datasets and analysis

After describing the results of their simulations, Martino used two published datasets as case studies to evaluate RPCA relative to Bray-Curtis and weighted UniFrac distances using PERMANOVA, KNN, and ordinations. The two datasets were drawn from a natural study comparing healthy and stressed sponges (1) and from a study that used a murine model of sleep apnea where mice were given air or intermittent hypoxia and hypercapnia (IHH) (2). The sponge dataset had 133 healthy and 85 stressed samples and the sleep apnea dataset had 96 air treated and 93 IHH treated samples (Figure 5). The effect sizes in both studies were relatively large and could easily be differentiated by the first two PCoA axes. Similar to the results from the simulations, I wondered whether differences between the methods would be observed when there was a smaller effect size. Therefore, I added a third dataset from a study that explored associations between the microbiome and colorectal cancer (3). The differences between healthy individuals and those with screen relevant colonic neoplasias were not obvious by PCoA and the study included more subjects. The colorectal cancer (CRC) dataset included 261 subjects with healthy colons and 229 subjects with screen relevant colonic neoplasias (Figure 5). The sponge, sleep apnea, and colorectal cancer datasets were analyzed in parallel.

**Figure 5.**
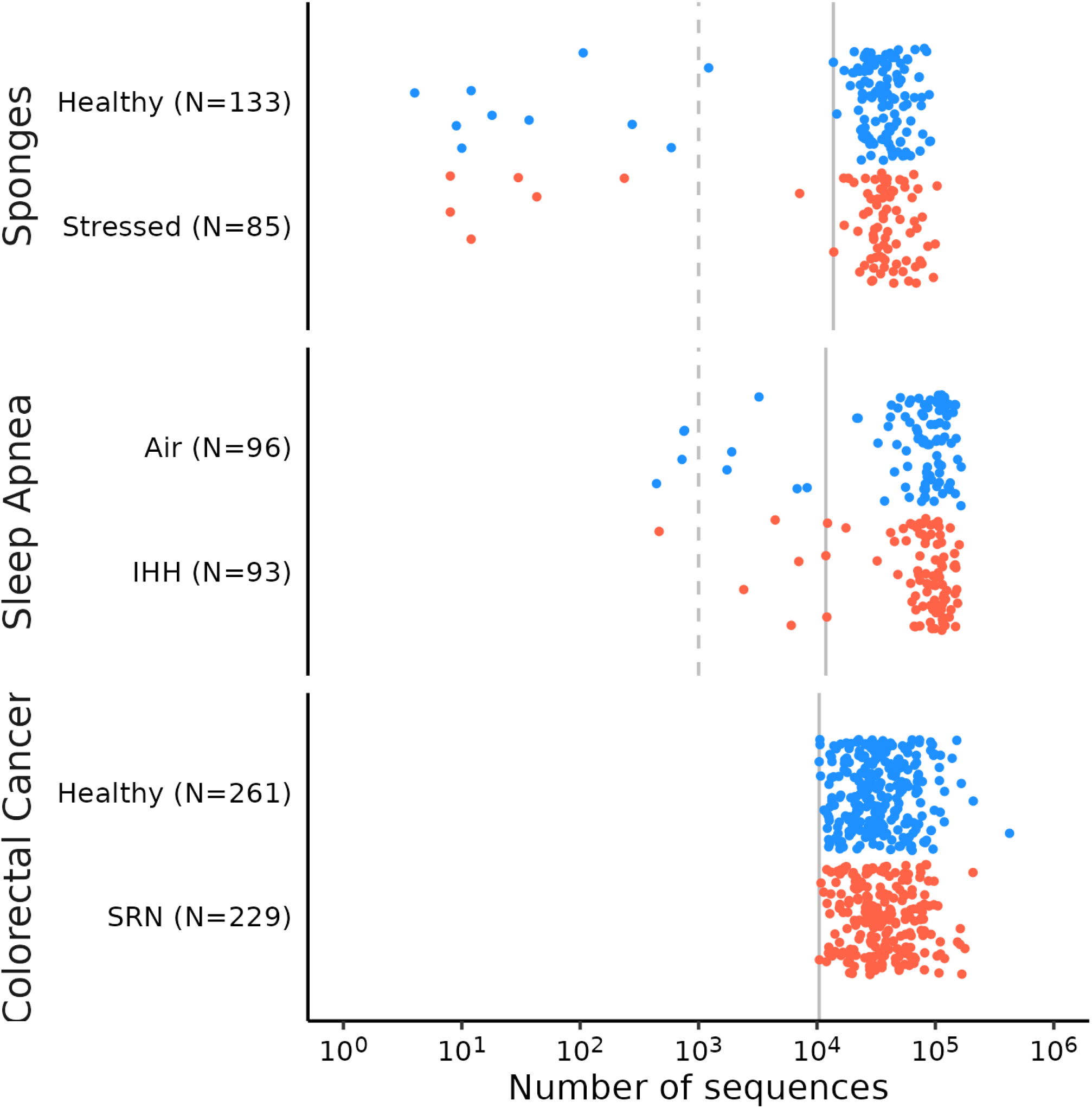
Martino selected an arbitrarily small number of sequences and a pair of datasets that were small relative to other available datasets. The vertical dashed line indicates the 1,000 sequence threshold, which was used by Martino and colleagues when calculating Bray-Curtis and weighted UniFrac distances. The vertical solid line indicates the higher threshold that was added in this study to explore the effects of sequencing depths. The higher threshold was 11,879 sequences for the sleep apnea dataset, 13,755 sequences for the sponges dataset, and 10,439 sequences for the colorectal cancer dataset.

After conducting an exhaustive review of the code that Martino and colleagues provided in their GitHub repository, I discovered several problems in the original analysis of the case studies. The details of this review can be found in the Supplementary Text. Aside from the selection of datasets with large effect sizes that were visible using the first two PCoA axes, the principle problems included (i) performing a single subsampling the samples in each data to an artificially low number of sequences (i.e., 1,000) for the Bray-Curtis and weighted UniFrac distance calculations; (ii) removal of sequences for the Bray-Curtis and weighted UniFrac distance calculations prior to subsampling that weren’t removed from the RPCA analysis because sequences were missing from the trees; (iii) they performed 9 rather than the 10 random subsamples of the datasets when generating balanced designs of the data sets (10 was described in the manuscript); (iv) running RPCA with a rank of 3 for the sleep apnea dataset rather than 2 as specified in the manuscript; (v) significant problems in the training and testing of the KNN classification to the data including the selection of the hyperparameter, lack of balance design in the stratifications, and only performing a single iteration; and (vi) only using two PCoA axes rather than the actual Bray-Curtis and weighted UniFrac distances in the KNN classification. The following analyses used the remedies as described in the Supplementary Text.

Similar to the analysis of the simulated datasets, pairwise Euclidean distances were calculated using 2 or 3 RPCA axes, Nudge, One, and Zero CLR transformed data, PCA and DeSeq2 transformed raw counts, raw counts, relative abundance data, single subsampling, rarefaction, and based on the first two PCoA axes using Bray-Curtis distances. Pairwise Bray-Curtis distances were calculated using raw counts, relative abundance data, SRS and CSS normalized data, single subsampling, and rarefaction. Pairwise weighted UniFrac distances were calculated using raw counts, SRS, single subsampling, and rarefaction; the calculation of weighted UniFrac distances includes converting raw counts to relative abundances (29).

### PERMANOVA analysis of case study datasets

I repeated Martino’s PERMANOVA analysis to test whether the average beta-diversity was significantly different between the two treatment groups within each of the three case studies. Their analysis varied the number of samples per treatment group to maintain a balanced design. This choice was peculiar since PERMANOVA is not sensitive to unbalanced sampling. When I used all of the samples available and performed the PERMANOVA analysis, the only dataset and methods that did not yield a significant result was for the colorectal cancer dataset using RPCA with 2 or 3 axes (P = 0.09 and 0.06, respectively). A second peculiar choice was to subsample the number of sequences to 1,000 sequences per sample when calculating Bray-Curtis distances but to use all of the sequences with RPCA. They could have increased the number of sequences by more than ten-fold without adversely impacting the number of samples in their analysis (Figure 5). Acceptable thresholds could have been 11,879 sequences for the sleep apnea dataset, 13,755 sequences for the sponge dataset, and 10,439 sequences for the colorectal cancer dataset. Overall, using the higher sequence count thresholds did not drastically alter the results of the analyses; however, it typically resulted in larger PERMANOVA F statistics (Figure 6). For the remainder of the PERMANOVA analysis, I have used the larger number of sequences instead of 1,000 sequences.

**Figure 6.**
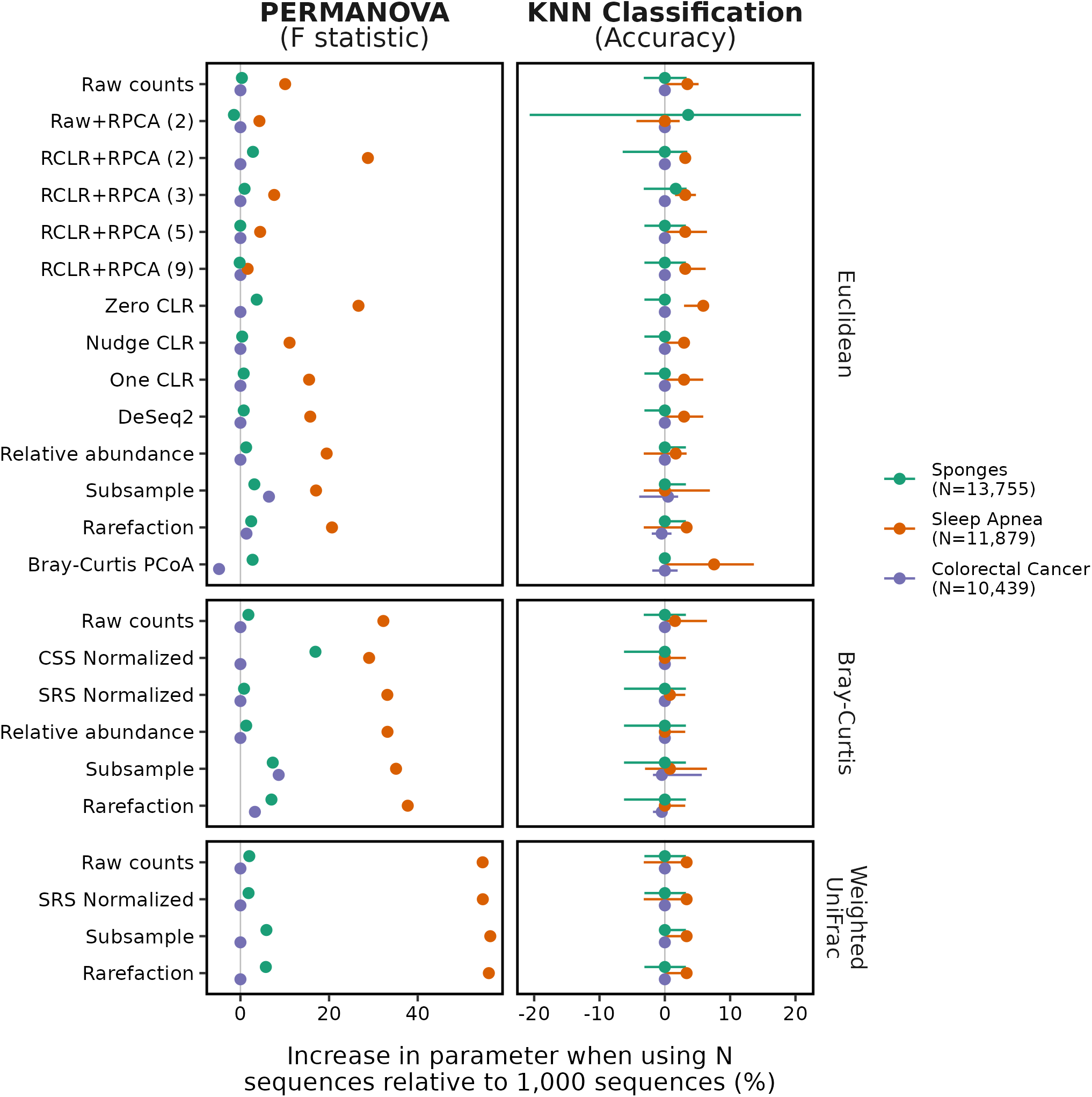
Increasing the minimum number of sequences per sample from 1,000 to more than 10,000 resulted in larger PERMANOVA F statistics and KNN accuracies. Calculation of the difference in F statistic was performed using values obtained when running PERMANOVA with all available samples. The difference in Bray-Curtis PCoA values was 220% and is not shown for the sleep apnea dataset because it was so much larger than the other values. Calculation of the difference in classification accuracy was performed using the largest possible balanced design, which included 78 samples for the sponges dataset, 87 samples for the sleep apnea dataset, and 229 samples for the colorectal cancer dataset. Because 10 iterations of the sample subsampling were performed we measured the difference in accuracy for each iteration. Each point represents the median difference and the confidence intervals represent the range. The vertical gray lines indicate where the F statistic or accuracy was not different between using 1,000 sequences and the number of sequences indicated for that dataset in the legend.

The results of my reanalysis of the sponge and sleep apnea datasets using PERMANOVA replicated what was observed in the original study (Figure 7). Across the various transformations and sample sizes, at least 9 of the 10 replicates yielded significant differences between the treatment groups. For the colorectal cancer dataset the results were more varied. The most striking result was that RPCA with 2 or 3 axes only yielded significant results in 2 or 3 of the 10 replicates. When 5 or 9 axes were used, the tests were significant more frequently. With the exception of Zero CLR and using Euclidean distances extracted from Bray-Curtis PCoA, the other transformations consistently yielded significant results. Across all methods, the fraction of significant tests increased with the number of samples in each group. The inclusion of the colorectal cancer dataset, which had a smaller effect size than the sponge and sleep apnea datasets helped to highlight the shortcomings of RPCA.

**Figure 7.**
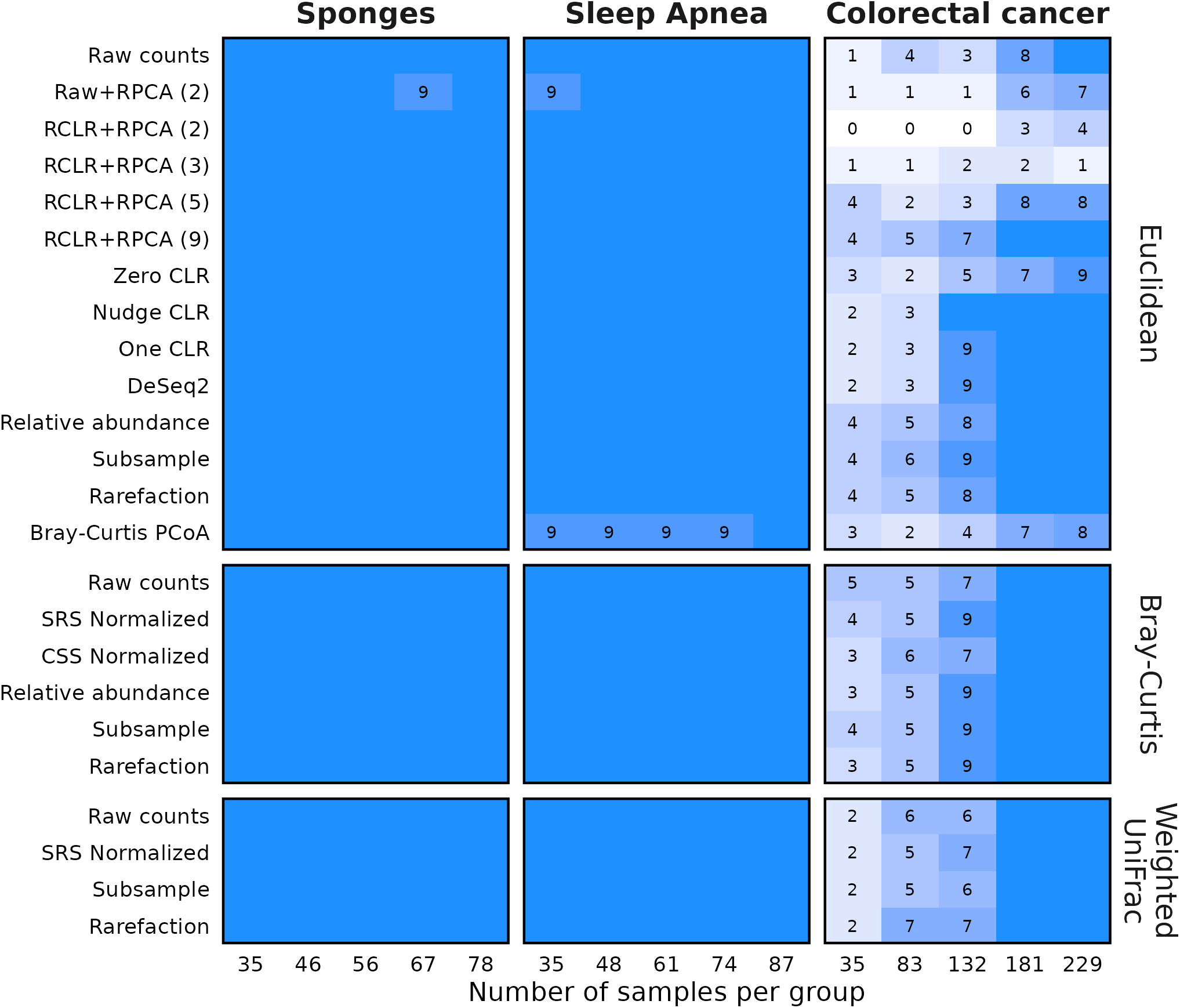
Although all methods perform well on datasets with a large effect size between treatment groups, the RPCA-based methods suffer relative to the others when the effect size is smaller. Each dataset was subsampled to the indicated number of samples per treatment group. For each of ten subsamplings, PERMANOVA was used to differentiate between the treatment groups. The numbers in each tile indicate the number of subsamplings that yielded a P-value less than 0.05. The blue tiles without numbers indicate those conditions where all 10 subsamplings yielded a significant result.

### KNN classification of case study datasets

I repeated Martino’s KNN-based analysis using the remedies described in the Supplementary Text on all three datasets with each of the methods for correcting for uneven sampling effort (Figure 8). Rather than using their strategy of using a fixed number of nearest neighbors, I repeatedly fit the hyperparameter for the number of neighbors (i.e., K) using 80:20 training and testing splits. Overall, using the higher sequence count threshold did not dramatically alter the results of the analyses; however, it typically resulted in higher accuracy to classify samples into the treatment groups for the sleep apnea dataset, which had fewer samples (Figure 6). For the remainder of this analysis, I will describe the results using the larger number of sequences.

**Figure 8.**
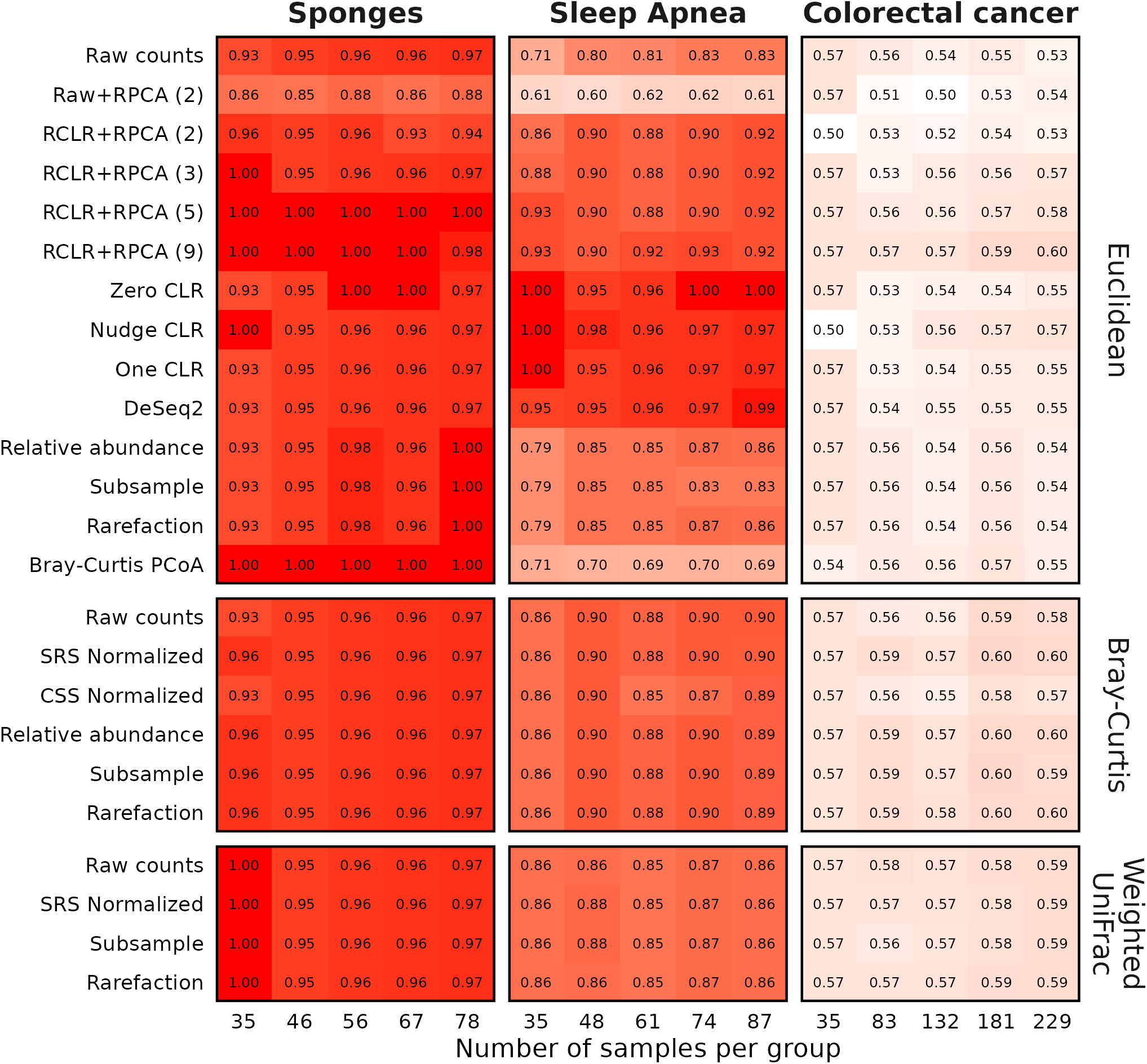
The RPCA-based methods performed worse or at best, no better, than other methods for classifying samples when using KNN classification. Each dataset was subsampled to the indicated number of samples per treatment group 10 times and was used to train and test a KNN classifier. The value in each cell is the median accuracy amongst those subsamplings.

Similar to the PERMANOVA analysis, my KNN classification results largely replicated those observed for the sleep apnea and sponge datasets by Martino. In general, using RPCA with any number of axes resulted in median accuracies greater than 0.90 for these datasets. When using RPCA with the colorectal cancer dataset the accuracy increased modestly with the number of axes and the effect of the number of samples per group did not have a consistent effect on classification accuracy. Caution should be used in interpreting such results as there was no *a priori* reason to use more than 2 axes. Within the sponge dataset there was little variation in the classification accuracy with most methods yielding values between 0.94 and 1.00; the worst performing method was using raw abundances as input to RPCA with two axes (accuracy = 0.88). With the sleep apnea dataset the CLR-based methods and DeSeq2 transformed data outperformed the RPCA methods, which outperformed the other Euclidean-distance methods; accuracies calculated using KNN with Bray-Curtis distances performed similar to the RPCA-based methods. Finally, with the colorectal cancer dataset, KNN classification using Bray-Curtis distances outperformed the Euclidean-based distances for all transformations except when RPCA was used with 5 or 9 axes. These results again demonstrate that RPCA-based methods performed worse than other methods when the effect size was small. Even for the sponge and sleep apnea datasets the differences in accuracy were negligible relative to other methods.

### Sampling depth experiment with simulated and experimental data

To test the sensitivity of various methods to uneven sequencing effort, Martino generated duplicated each of the case study samples by subsampling one copy to 500 sequences per sample and subsampling the other copy to 100 sequences per sample. They calculated distances using Jaccard, Bray-Curtis, and RPCA and then visualized the distances using ordination. Martino stated that “RPCA alleviated the clustering by sequencing depth both qualitatively and quantitatively”. However, in the analysis presented in their Supplementary Materials, the ordinations still showed clustering by sampling depth for RPCA. Furthermore, the Jaccard and Bray-Curtis distances were not calculated using rarefaction or even a single subsampling, which are the typical strategies to control for such uneven sampling with these metrics. To reassess the sensitivity of RPCA and other methods to uneven sequencing effort, I repeated their analysis with the simulated datasets generated using the original and revised code and the case study datasets. Instead of using 100 and 500 sequences per sample, I used 5,000 and 10,000 sequences per sample. A two factor PERMANOVA design (i.e., depth and treatment group) was used to test for significant differences in the level of sequencing effort. The results of this analysis were consistent with other analyses in this and other studies (Figure 9). Raw counts, relative abundances, RPCA-based distances, CLR-transformed counts, DeSeq2, and CSS normalized data yielded significant differences between the number of sequences per sample. Those methods were not able to control for uneven sampling effort. In contrast, SRA normalization, subsampling, and rarefaction consistently controlled for the effects of uneven sampling effort across the simulations.

**Figure 9.**
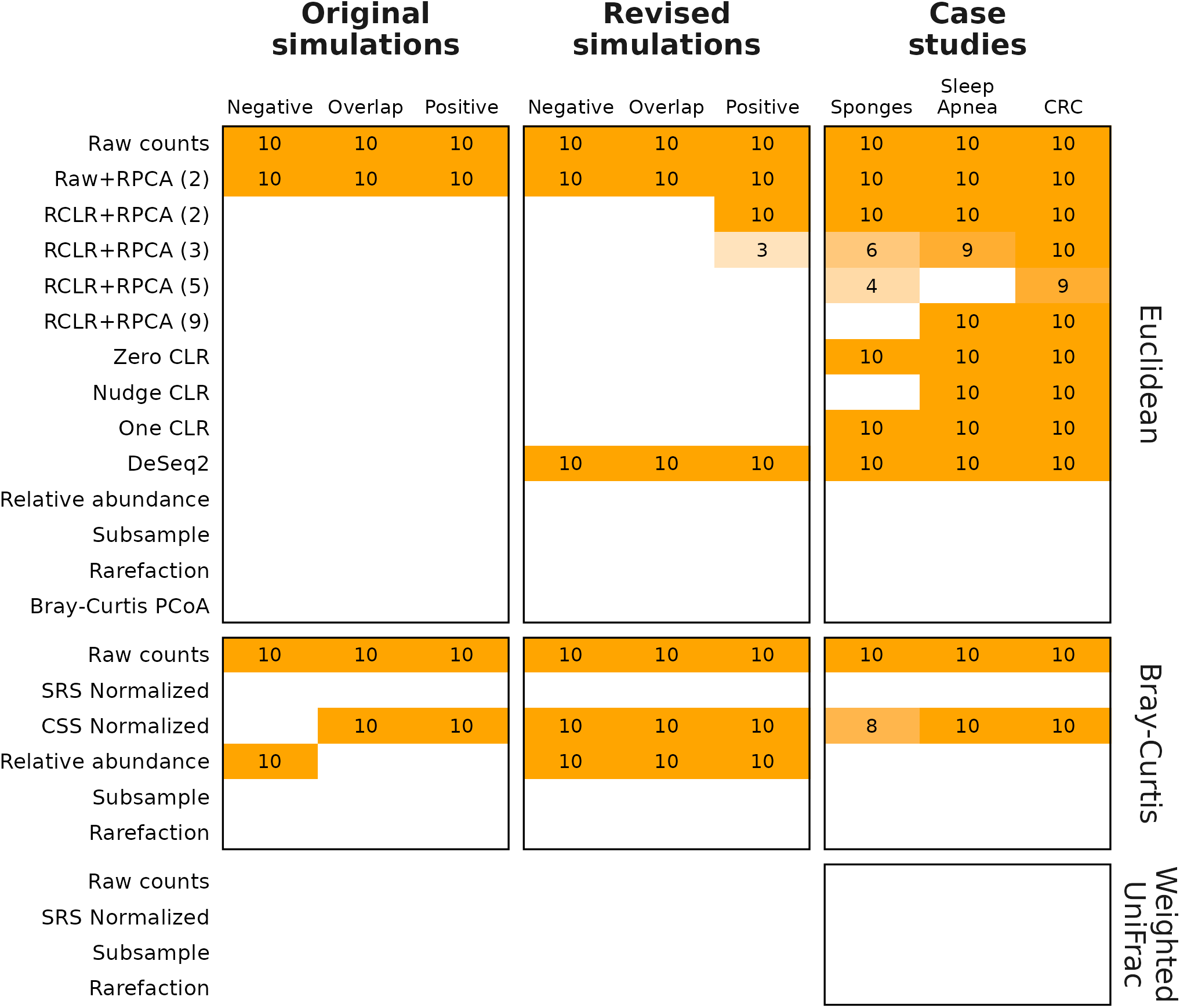
RPCA did not alleviate clustering of samples by sequencing depth; rarefaction and subsampling did. The numbers within each cell indicate the number of simulations out of 10 that falsely gave a significant P-value for the comparison of sequencing depth. White cells without a number had no falsely significant P-values. Data are not provided for the simulations analyzed with Weighted UniFrac distances since there were no trees. Synthetic versions of each dataset were generated where each sample was subsampled to 5,000 and 10,000 sequences, combined, processed, and analyzed by a two-factor PERMANOVA. Each simulation was repeated 10 times.

## Discussion

When evaluating bioinformatics tools, simulated datasets are useful when a null or true difference between treatment groups in a real dataset cannot be established. The simulation approach described by Martino offered a novel approach to studying the performance of various methods for controlling uneven sampling effort. Unfortunately, as implemented in the original study, the simulation code had critical errors that made it impossible to meaningfully compare overlapping treatment groups. When the code was corrected and a smaller effect size was applied, the limitations of RPCA relative to other methods became clear. Because simulated data does not not always depict the complexity of real data, real datasets can be useful for demonstrating the utility of these methods. The datasets selected by Martino had large effect sizes, which limited the ability to detect variation in performance since the differences were relatively easy to detect. For example, significant P-values and median classification accuracies above 0.90 were common regardless of the method. By incorporating the colorectal cancer dataset, which had a small effect size, there was more variation in the power to detect differences and classification accuracy. Under more nuanced conditions RPCA was no better than rarefaction and was often worse. Since a researcher is unlikely to know the effect size before conducting their analysis, it would be most prudent to select methods with the most power.

Martino implemented methods for evaluating RPCA relative to other strategies of controlling for uneven sampling effort that would not normally be used in a typical microbiome analysis. First, Martino benchmarked RPCA by comparing the method’s performance using raw counts or RCLR-transformed data. Missing from their benchmarking was a comparison to Bray-Curtis or UniFrac distances or use of other methods of controlling for uneven sampling effort. The result was a comparison that made their new method appear considerably better than a method that no one else has used in a published microbiome analysis. Second, they used KNN as a classification tool rather than using logistic regression or random forest. KNN is simply not used for classifying microbiome data. Compounding the problem of using KNN was that they did so based on distances from the first two or three axes of a PCoA using a Bray-Curtis or weighted UniFrac distance matrices rather than directly using the ecological distances themselves. The effect was a loss of information stored in higher dimensions. Even with the limitations of KNN (e.g., requirement of balanced design), the other methods outperformed RPCA.

There are several limitations inherent in RPCA that would still limit its use even if its performance were better. First, there is no guidance on selecting the number of axes one should use with RPCA. Martino used 2 axes with their simulated and sponges dataset and 3 axes with the sleep apnea dataset; 3 is the default value in their publicly released gemelli software for running RPCA. As I showed with the real datasets, increasing the number of axes increased the power to detect differences and the classification accuracy. This creates a scenario where users could select a number of axes to fit their expected results (i.e., “P-hacking”). Second, as Martino indicated the method does not perform well with time series data or where there is considerable variation between the samples. This is likely when effect sizes between treatment groups are small as is typical in most microbiome analyses.

Martino made the claim in their paper’s Discussion section that Aitchison distances calculated using RPCA are insensitive to uneven sequencing effort. The only data they provide to support this claim was found in their paper’s Materials and Methods section, which references their Supplemental Figure 3 and Supplemental Table 2. Yet, those data show that the RPCA ordination also clustered by sequencing depth. Unfortunately, they did not rarefy the Jaccard or Bray-Curtis distances to a common number of sequences as one would do in a real analysis. I replicated this analysis, but with a larger number of sequences per sample using simulated and real datasets, and using a two-factor PERMANOVA analysis. These results (Figure 9) confirmed that RPCA, the other CLR-based methods, using raw counts, DeSeq2, CSS normalization, and relative abundance were unable to control for uneven sampling effort in this analysis. Only rarefaction, subsampling, and SRS normalization were able to control for uneven sampling effort.

This is my third analysis describing the effects of uneven sequencing effort and comparing the methods to control for those negative effects (4, 25). My results in this study confirm what I previously reported. The primary difference is that subsampling and SRS normalization performed as well as rarefaction. It is worth noting that in a previous analysis where I correlated sequencing depth with multiple alpha and beta diversity metrics the correlations for SRS normalization varied widely across 12 different real datasets. The benefit of rarefaction over subsampling is that distances are based on a large number of iterations (e.g. 100 or 1,000) rather than only one. Until it is possible to obtain a uniform number of sequence reads per sample, rarefaction should be used to control for uneven sampling effort.

## Materials & Methods

### Deicode benchmarking data and code

Martino made a git-based repository available on GitHub that includes the actual benchmarking code written in Python and the data used in their analysis (https://github.com/cameronmartino/deicode-benchmarking). Deicode was the original name of the package that implemented RPCA. That functionality is now available as part of the gemelli package. The code and data from their benchmarking repository are included as the directory “deicode_benchmarking” within repository for the current analysis (https://github.com/SchlossLab/Schloss_RPCA_mSystems_2026). A conda virtual environment was created to mimic the versions of the tools used the original simulation. The environment file is available within the current project’s repository as workflow/envs/rpca.yml

### Revised simulation code

A revised version of Martino’s simulation code was written in R as described in the Results section. The code I generated used functions from base R and the tidyverse metapackage. The script that implemented the simulation is available in the repository as workflow/scripts/simulations.R. Similar to Martino’s original Python implementation, the simulation code took as input the number of samples, OTUs, treatment groups, OTUs shared between the treatment groups, and sequences to generate the simulated communities. A driver script subsampled the data to the desired number of sequences and formatted the data for downstream analysis.

### Case study data

The operational taxonomic unit (OTU) count data and metadata for the sponges and sleep apnea datasets were obtained from the Deicode benchmarking repository. Because the trees that were included with the benchmarking repository did not contain all of the sequences contained within the OTU tables, I generated new trees. The OTU identifiers in the count data were the DNA sequences. I aligned the sequences to the NR v138.2 SILVA reference alignment (30, 31) and calculated pairwise distances between sequences (32). These distances were used as input to generate traditional neighbor joining trees (33). The curated sequence data and metadata for the colorectal cancer dataset were obtained from a previous study where it was referred to as the “Human” dataset (4). Those data were generated by obtaining sequence data and metadata from the Sequence Read Archive. The sequence data was processed using the mothur software program (34). This included assembling the paired reads that covered the V4 region of the 16S rRNA gene using a quality score-aware method (35), aligning the assembled contig to a SILVA reference alignment (30, 31), pre-clustering to remove residual sequencing errors (36) chimera checking and removal (37), and clustering of sequences into OTUs using the OptiClust algorithm (38). OTUs were defined a group of sequences not more than than 3% different from each other. Representative sequences from each OTU were used to generate traditional neighbor joining trees as described for the sponges and sleep apnea datasets.

### Distance-based K-nearest neighbor (KNN) classification

Martino’s KNN classification analysis used the KNeighborsClassifier function from scikit-learn (39). Although that function could have taken a pre-computed distance matrix as input in the version of scikit-learn that Martino used (v.0.5.9), they gave the function the ordination results with the desired number of axes. I reimplemented an R-based version of the KNN classification method as workflow/scripts/test_train_knn.R. This method implemented the training-testing framework described by Topcuoglu et al. (28) to fit the number of neighbors hyperparameter (i.e., K). My R script took as input a distance matrix, a design file that assigned samples to treatment groups, the fractions of data to be assigned to the training and testing groups, the number of inner and outer loops, and the hyperparameter grid for fitting K.

### Methods of controlling for uneven sampling effort

I evaluated twelve methods for controlling for uneven sampling effort. These included using: (i) raw counts, which were the number of sequences in each OTU; (ii) the OptSpace algorithm performed on raw counts as implemented in Deicode (v.0.1.5) (23); (iii) the full RPCA algorithm performed using gemelli (v.0.0.9) with 2, 3, 5, and 9 components (23); (iv) Zero CLR, which calculated the center log ratio (CLR) after imputing the of zero counts using the zCompositions (v.1.4.0.1) R package (20); (v) Nudge CLR, which calculated the CLR after adding a pseudocount of 1 divided by the total number of sequences in a sample to the abundance of each OTU in the sample (21, 22); (vi) One CLR, which calculated the CLR after adding a pseudocount of 1 to the abundance of all of the OTUs (21, 22); (vii) the DESeq2 R package (v.1.34.0), which was used to implement the variance stabilizing transformation to normalize the OTU counts (8, 14); (viii) the SRS R package (v.0.2.3), which was used to normalize OTU counts using the scaled ranked subsampling algorithm (12); (ix) the metagenomeSeq R package (v.1.36.0), which was used to normalize OTU accounts using the cumulative sum scaling (CSS) algorithm (13); (x) relative abundances, which were calculated as the abundance of each OTU in a sample divided by the total number of sequences in the sample; (xi) subsampling, which involved randomly selecting a specified number of sequences from each sample and removing any samples that had fewer than the desired number of sequences; and (xii) rarefaction, which involved repeating subsampling 100 times and calculating a distance matrix after each iteration followed by calculating the mean distance matrix.

### Methods for calculating distances between communities

The vegan R package (v.2.6_4) was used to calculate Euclidean and Bray-Curtis distance matrices (40, 41). Euclidean distances are not typically used in microbiome analyses because they inflate the similarity of communities. However, they are used with CLR transformed data resulting in so-called Aitchison distances. The OptSpace and RPCA algorithms outputted Euclidean distances directly from the Deicode and gemelli software. Euclidean distances were calculated for raw counts, and data transformed by the CLR-based methods, DeSeq2, relative abundance, subsampling, and rarefaction using the vegdist function in vegan. Because OptSpace, CLR-based methods, and DeSeq2 outputted negative values, it was not possible to calculate Bray-Curtis distances on those data. Bray-Curtis distances were calculated using raw counts, CSS and SRS normalized counts, relative abundances, and subsampled data using the vegdist function in vegan; rarefaction was performed using vegan’s avgdist function. The mothur package (v.1.48.5) was used to generate weighted UniFrac distances using raw counts, SRS normalized counts, subsampled data, and rarefaction (29); the mothur version produced the same distances as those from qiime2 using raw counts, but facilitates the calculation of distances using subsampling and rarefaction. The weighted UniFrac algorithm calculates relative abundances from the inputted counts and is not amenable to non-count data such as CSS normalized counts.

### Measuring effects of controlling for uneven sampling effort on performance of PERMANOVA

I used the adonis2 function from the vegan R package (v.2.6_4) to calculate the PERMANOVA F statistic for comparisons of treatment groups in the simulations and case studies (40, 42). Euclidean or Bray-Curtis distance matrices and a map indicating which group each sample belonged to were used as input to the function. Except for the simulation measuring the ability of the various methods to control the effects of sampling depth a single classification design was used in adonis2 to test the model, Distance ∼ Treatment

### Measuring ability of various methods to limit the effects of sampling depth

Following the approach described by Martino to assess the sensitivity of the different methods to uneven sequencing I randomly sampled 5,000 and 10,000 sequences from each of the simulation and case study datasets. These data were pooled so that each sample was represented at each of the sequencing levels. A sequencing depth variable was added to the treatment group label. Each dataset was replicated 10 times with a different random number generator seed. To test the effects of sequencing depth, I used the adonis2 function from the vegan R package (v.2.6_4) (40) to test the model, Distance ∼ Treatment + Sequencing Depth + Treatment x Sequencing Depth. Although this model tested for a treatment effect, it’s P-value was biased downward. This was because every sample was artificially duplicated, which made each treatment look more similar to itself than it really was, leading to reduced the intra-treatment variation. The result was a larger F statistic and smaller P-value for the treatment effect. My analysis of this simulation only focused on the significance of the Sequencing Depth effect.

## Data availability

No new biological data were generated in this study. The entire codebase for my analysis is available as a GitHub repository at https://github.com/SchlossLab/Schloss_RPCA_mSystems_2026. The repository includes a copy of Martino’s benchmarking repository. My repository includes a Snakemake-based workflow (v.7.32.4) includes all of the steps in the analyses described in this manuscript. The workflow uses R (v.4.3.3) and Python (v.3.11.14) packages and code. Notable R packages not already indicated included tidyverse (v.2.0.0), data.table (v.1.14.8), ggh4x (v.0.2.6), ggtext (v.0.1.2), patchwork (v.1.1.3), quarto (v.1.4.550), and reticulate (v.1.43.0). Notable Python packages not already indicated included matplotlib (v.3.8.1), numpy (v.1.12.1), pandas (v.2.1.3), scikit-bio (v.0.5.9), scikit-learn (v.1.3.2), and scipy (v.1.10.0).

## Acknowledgements

I am grateful to the reviewers of the papers describing my previous analyses who raised the question of why I saw different results from other developers who had developed methods of controlling for uneven sampling effort. I appreciate that Martino and colleagues made their benchmarking code and data publicly accessible. This study would not have been possibly without their commendable transparency. No generative artificial intelligence tools were intentionally used in this study or in the preparation of this manuscript.

## Supplemental Figures

**Figure S1.**
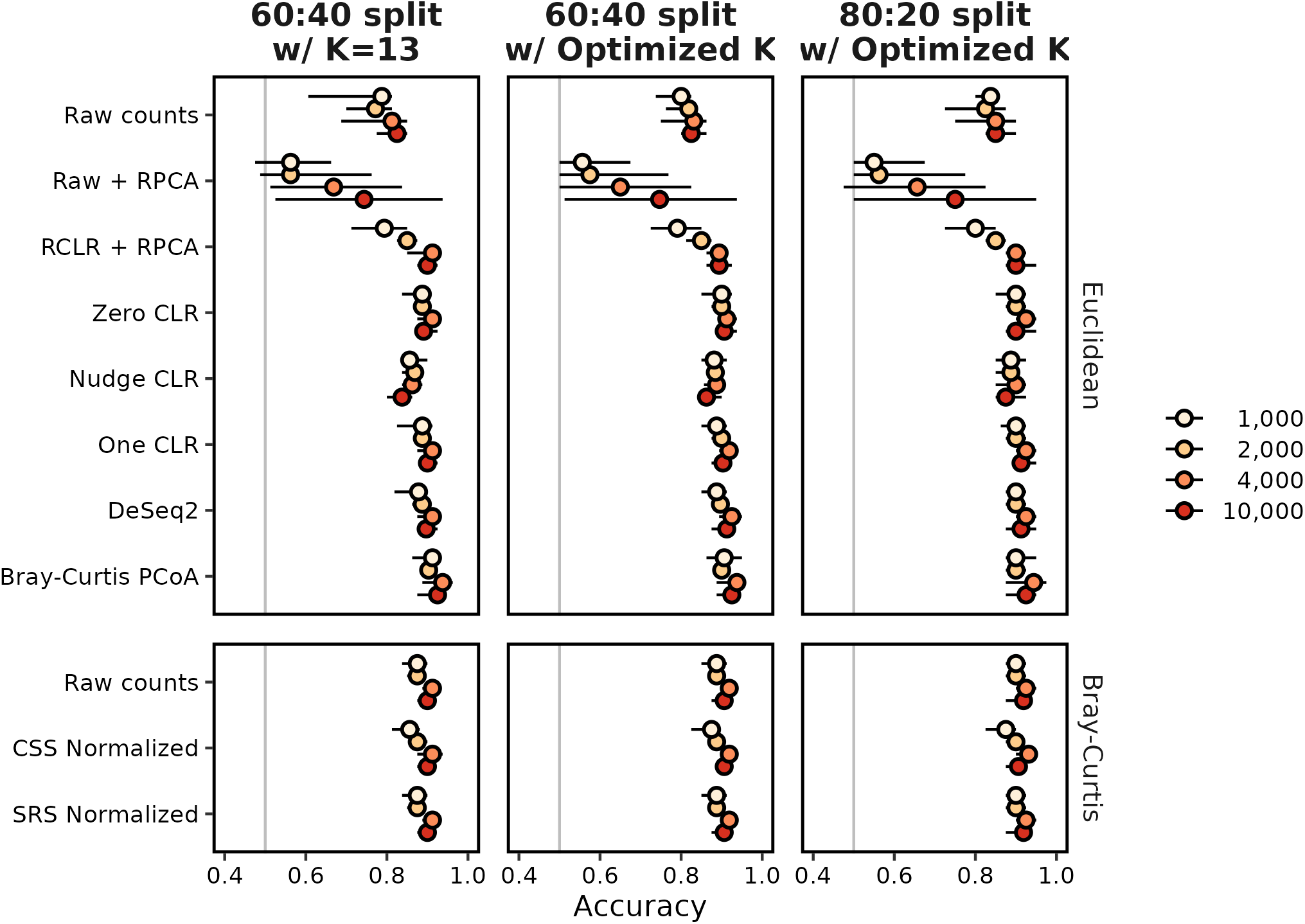
The choice of K and the size of the training and testing data sets had a minimal effect on KNN classification accuracy when using the original Martino overlap simulation data. Martino split 60% of their data into a training set using 13 neighbors and tested on the remaining 40%. Following Topcuoglu et al. (28), I optimized the number of neighbors using 60:40 and 80:20 training-testing splits. The 80:20 data were used in the Results section.

**Figure S2.**
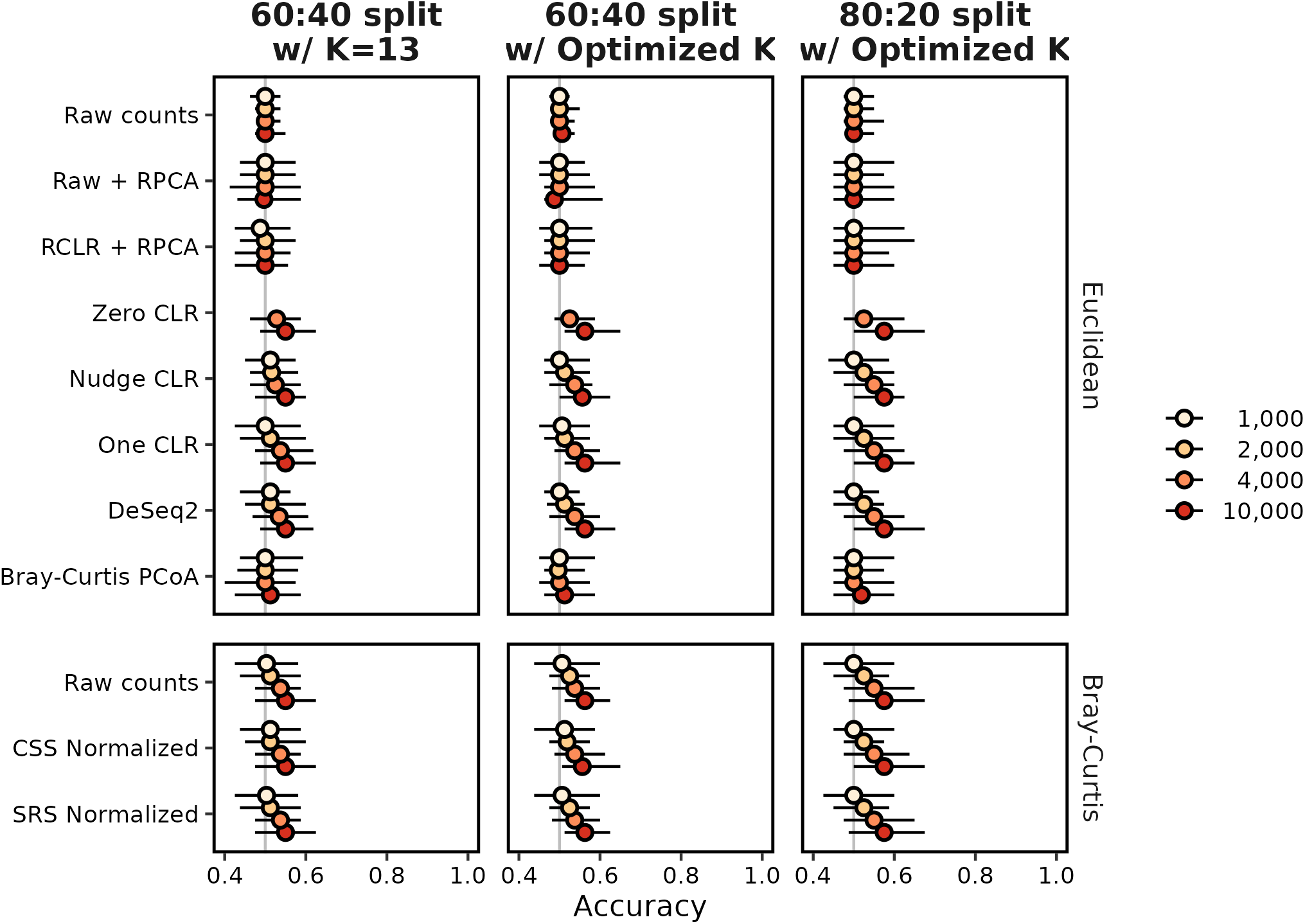
The choice of K and the size of the training and testing data sets had a minimal effect on KNN classification accuracy when using the data generated by the revised simulation code. Martino split 60% of their data into a training set using 13 neighbors and tested on the remaining 40%. Following Topcuoglu et al. (28), I optimized the number of neighbors using 60:40 and 80:20 training-testing splits. The 80:20 data were used in the Results section.

## Notes

### Competing Interest Statement

The authors have declared no competing interest.

### Summary of Updates

Modified to address peer review at mSystems

