## Supplementary Text for "Rarefaction is better than robust Aitchison PCA and other compositional data analysis methods at controlling for uneven sequencing effort"

### Supplemental text: Review of Python code from Martino et al.

**Review of code used to generate positive and negative control heatmaps in Figure 3DE of Martino** **1**

**Review of build_block_model function and its dependencies** **3**

**Literature cited** **23**

The code chunks displayed below were copied directly from the code used by Martino et al. (1) and the line numbers correspond to the indicated files. The hyperlinks in the text go to the GitHub repository for the current project, which was a copy of the original repository developed by Martino. Martino's repository was forked from the Deicode repository on January 28, 2019. The only changes between the Martino repository and the copy for this project was to make the directory and file names lowercase and to replace hyphens with underscores.

#### Review of code used to generate positive and negative control heatmaps in Figure 3DE of Martino

Review of the code in `deicode_benchmarking/simulations/negative_control.ipynb` revealed two important problems with the generation of Martino's Figure 3DE. For background, the `X_random` and `X_signal` matrices represented the negative and positive controls, respectively, that were generated in an earlier block of this notebook. For both matrices, there were 1000 rows for each feature or taxon and 200 columns for each sample. At L13 and L15 of block 183, the `imshow` function from the pyplot package within the `matplotlib` Python library was used to generate heatmaps of the two matrices.

```
10 #show the results
11 fig,((ax1,ax2),(ax3,ax4)) = plt.subplots(2,2,figsize=(12,8))
12 
13 ax1.imshow(clr(X_signal+1),aspect='auto',norm=MidpointNormalize(midpoint=0.), cmap='PiYG')
14 ax1.set_title('Positive Control Simulation',fontsize=22)
15 cbim = ax2.imshow(clr(X_random+1),aspect='auto',norm=MidpointNormalize(midpoint=0.), cmap='PiYG')
```

```
16 ax2.set_title('Negative Control Simulation',fontsize=22)
```

Within the arguments to `imshow` (L13), the matrices were transformed by adding 1 to each cell and then subjecting the matrices to a central log ratio transformation with the `clr` function from the composition statistics module of the `skbio` Python library. It is important to note that the `clr` function assumed that the rows represented compositions or samples and the columns represented the components or taxa. Therefore, `X_random` and `X_signal` should have been transposed prior to performing the clr transformation. For example, earlier at L2 and L6 of the same code block, the matrices were transposed prior to using the `rclr` function.

```
1 #RPCA on random
2 X_random_rclr = rclr().fit_transform(X_random.T)
3 opt_noise = OptSpace().fit(X_random_rclr)
4 U_random = pd.DataFrame(opt_noise.sample_weights,index=X_random.columns)
5 #RPCA on very clear signal
6 X_signal_rclr = rclr().fit_transform(X_signal.T)
7 opt_sig = OptSpace().fit(X_signal_rclr)
8 U_signal = pd.DataFrame(opt_sig.sample_weights,index=X_signal.columns)
```

As executed in the notebook, the result of using the `clr` function was the same shape as the input matrices with features as rows and samples as columns. The transformed data were then used in the `imshow` function which plotted the rows and columns of the matrix to the rows and columns of the heatmap. However, at L21-24 in the same code block, "Samples" was added to the y-axis label and "Features" was added to the x-axis label, which was the reverse of the actual situation. These problems hid the fact that samples rather than taxa were shared between the treatment groups. Of course, sharing samples between treatment groups would not make sense and instead the taxa should have been shared.

```
21 ax2.set_ylabel('Samples',fontsize=15)
22 ax1.set_ylabel('Samples',fontsize=15)
23 ax2.set_xlabel('Features',fontsize=15)
24 ax1.set_xlabel('Features',fontsize=15)
```

The problem with the labelling of the axes could be more clearly seen by changing the value of the `aspect` argument in `imshow` from `'auto'` to `None` on L13-14. To summarize, the code in this block (i) performed the CLR transformation across the samples rather than across the taxa and (ii) mislabelled the axes of the heatmaps in Martino's Figure 3DE. Both problems could have been avoided by transposing the matrices within the arguments to `clr` at L13 and L15. The comparison between transposing the data or not are are shown below.

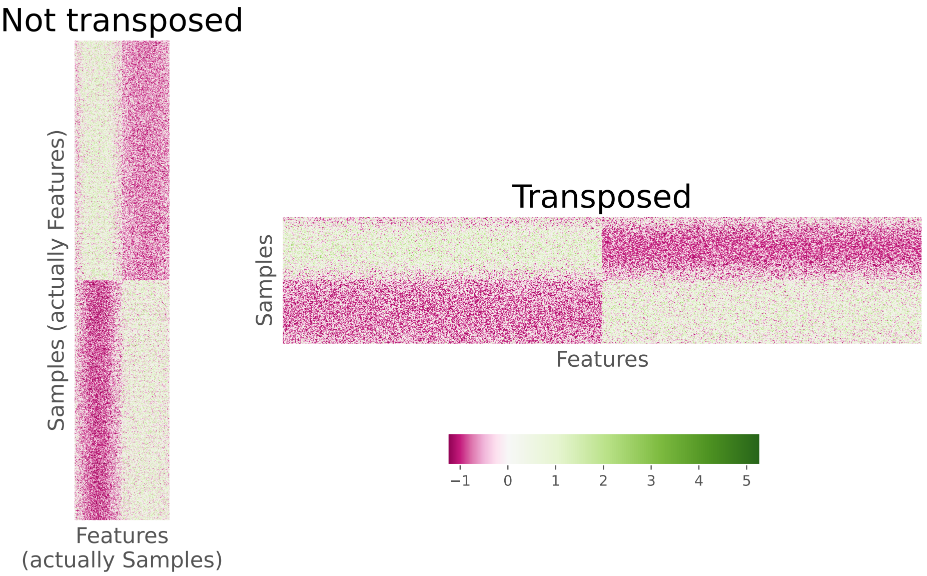

#### Review of `build_block_model` function and its dependencies

Because it was clear that there was a problem in the simulation code such that samples rather than taxa were shared between treatment groups, I reviewed the code in `build_block_model` and related functions to better diagnose the problems in the simulation. The `build_block_model` function was found in `deicode_benchmarking/simulations/scripts/simulations.py`. The function was called in `deicode_benchmarking/simulations/negative_control.ipynb` to generate the positive control data in Martino's Figure 3D and in `deicode_benchmarking/simulations/scripts/cluster_build.py` to generate the overlapping data analyzed in Martino's Figure 3ABC. The `build_block_model` function is listed below except for lines corresponding to comments (L185-236) and code that did not impact the generation of `X_noise` (L244-253 and L262-267). The function returned the values of `X_true` and `X_noise`.

```
175 def build_block_model(
176         rank,
177         hoced,
178         hsced,
179         spar,
180         C_,
181         num_samples,
182         num_features,
183         overlap=0,
184         mapping_on=True):
```

```
237     X_true = block_diagonal_gaus(
238       num_samples,
239       num_features,
240       rank,
241       overlap,
242       minval=.01,
243       maxval=C_)
```

```
254     X_noise = X_true.copy()
255     X_noise = np.array(X_noise)
256     # add Homoscedastic noise
257     X_noise = Homoscedastic(X_noise, hoced)
258     # add Heteroscedastic noise
259     X_noise = Heteroscedastic(X_noise, hsced)
260     # Induce low-density into the matrix
261     X_noise = Subsample(X_noise, spar, num_samples)
```

As called in `cluster_build.py`, the value of `rank` was 2 (i.e., the number of treatment groups), `num_samples` was 200 (i.e., the number of samples across all treatment groups), `num_features` was 1000 (i.e., number of taxa or features), and `overlap` was 20 (i.e., the number of features that overlapped between the treatment groups; it was zero in the positive control analysis). The values of `spar`, `C_`, `hoced`, and `hsced` were based on a parameter that was altered for each sequencing depth. The `spar` and `C_` values were set to the same value (e.g., 2500) and were used to scale the relative abundance values in the `Subsample` (L261) and `block_diagonal_gaus` (L237-243) functions, respectively. As used in the simulations with partially overlapping features the `hoced` and `hsced` values were set to `hoced` and `hsced` values divided by 15; for the positive control analysis they were divided by 60. These values were used in the `Homoscedastic` (L257) and `Heteroscedastic` (L259) functions where `hoced` and `hsced` were used to set the standard deviation of a normal distribution with mean set to the relative abundance of the sample-taxon pair receiving additional noise uniformly or randomly. The values of `spar`, `C_`, `hoced`, and `hsced` used in `build_block_model` were selected by fitting the model to experimental data from a study comparing the microbial communities on subject's fingers and keyboards. Although I did not attempt to replicate their model fitting, other code was found in `simulations.py` that ostensibly performed this analysis.

I noticed that when I loaded `simulations.py` and ran `build_block_model` multiple times with the same input data and random number generator seed, the code did not produce the same results. Although not a critical problem, the lack of computational reproducibility was part of my motivation to assess the benchmarking test results obtained with ten rather than only one seed.

Finally, comments in `build_block_model` described the function as returning a data frame with `num_samples` rows and `num_features` columns (L223-227). In fact, it returned a data frame with `num_features` rows and `num_samples` columns. This may account for why samples rather than features overlapped in the `block_diagonal_gaus` function. This issue is described further in the next section.

```
223     Returns
224     -------
225     Pandas Dataframes
226     Table with a block diagonal where the rows represent samples
227     and the columns represent features.  The values within the blocks
228     are gaussian.
```

#### Review of `block_diagonal_gaus` function

The function `block_diagonal_gaus` created a model that described the abundance of each taxon across the samples for each treatment group before any noise was applied to the data. This function was called by `build_block_model` and was also found in `deicode_benchmarking/simulations/scripts/simulations.py`.

```
def block_diagonal_gaus(
        ncols,
        nrows,
        nblocks,
        overlap=0,
        minval=0,
        maxval=1.0):
```

As used in Martino, `ncols` was 200 and `nrows` was 1000. As described in the Methods of the original study, the model used a normal distribution to vary the relative abundance of each taxa with different distributions for each treatment group such that a specified fraction of taxa overlapped between the two treatment groups. Within `block_diagonal_gaus` the number of treatment groups was specified with `nblocks`, which was 2 and the overlap was specified with `overlap`, which was 20 for the partial overlap simulation and 0 for the positive control simulation. The value of `maxval` was used to scale relative abundance values in the function and was set to the parameter described in the previous section (e.g., 2500 for a sequence depth of 4000). The function generated a matrix with `ncols` samples and `nrows` taxa; however, comments in the function described a transposed version of the matrix. In addition, as I'll show below, the output was not a block diagonal.

```
    Returns
    -------
    np.array
        Table with a block diagonal where the rows represent samples
        and the columns represent features.  The values within the blocks
        are gaussian distributed between 0 and 1.
```

```
117     if nblocks <= 1:
118         raise ValueError('`nblocks` needs to be greater than 1.')
119     mat = np.zeros((nrows, ncols))
120     gradient = np.linspace(0, 10, nrows)
121     mu = np.linspace(0, 10, ncols)
122     sigma = 1
123     xs = [norm.pdf(gradient, loc=mu[i], scale=sigma)
124           for i in range(len(mu))]
125     mat = np.vstack(xs).T
```

The function began by initializing a matrix with zeroes (i.e., `mat`) and a gradient of values to define a distribution (i.e., `gradient`). However, this code was largely superfluous since the values of these parameters were written over by subsequent code before the matrix (i.e., `mat`) was updated. Regardless, operationally, `mat` was defined as a matrix of zeroes with `num_features` rows and `num_samples` columns (L119). A vector with `num_features` values, `gradient`, was defined as a linear interpolation between the values of 0 and 10 (L120). Similarly, a vector with `num_samples` values, `mu`, was defined as a linear interpolation between the values of 0 and 10 (L121). These values along with a `sigma` value of 1 were used to create a `num_samples` long vector of arrays, which were each `num_features` long (L123-124). This vector of arrays, `xs`, took its values from the Normal probability density function (i.e., `norm.pdf`) at the `num_features` values of `gradient` under a distribution with a mean at each of the `num_samples` values of `mu` with a standard deviation of `sigma` (i.e., 1). The `num_features` by `num_samples` matrix `mat` was generated by combining each element of `xs` as the columns. As suggested in the comment above from L108-110, this matrix had maximum probabilities along the diagonal of the matrix. Values far off the diagonal were nearly zero. For example, the value for the first feature in the first sample was 0.3989 and it was 7.7e-23 in the last sample. The following heatmap shows the raw values in the matrix (the matrix has been transposed).

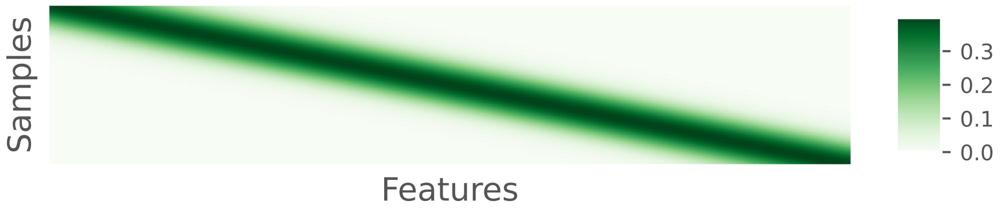

As stated above in my description of L117-125, the values of `mat` were largely overwritten by the following lines of code.

```
127     block_cols = ncols // nblocks
128     block_rows = nrows // nblocks
129     for b in range(nblocks - 1):
```

```
130
131         gradient = np.linspace(5, 5, block_rows)  # samples (bock_rows)
132         # features (block_cols+overlap)
133         mu = np.linspace(0, 10, block_cols + overlap)
134         sigma = 2.0
135         xs = [norm.pdf(gradient, loc=mu[i], scale=sigma)
136               for i in range(len(mu))]
137
138         B = np.vstack(xs).T * maxval
139                 lower_row = block_rows * b
140                 upper_row = min(block_rows * (b + 1), nrows)
141                 lower_col = block_cols * b
142                 upper_col = min(block_cols * (b + 1), ncols)
143
144                 if b == 0:
145                     mat[lower_row:upper_row, lower_col:int(upper_col + overlap)] = B
146                 else:
147                     ov_tmp = int(overlap / 2)
148                     if (B.shape) == (mat[lower_row:upper_row,
```

L127 and L128 defined the number of columns (`block_cols`) and rows (`block_rows`) in each block. Because `nblocks` was 2 for all simulations, these values became 100 and 500, respectively. At L129, a loop was initiated for values of an index variable, `b`, which ranged between 0 and the integer less than `nblocks-1` or 0. Therefore, the loop was only executed once and the else statement at L146-159 was never executed. At L131 and L133 new versions of `gradient` and `mu` were generated. All the values in `gradient` were 5 and the values in `mu` were taken from a linear extrapolation between 0 and 10. The vector `mu` had `block_cols` (i.e., 100) plus `overlap` (i.e., 20) values. Because the columns contained sample data, the additional 20 columns represented 20 additional samples rather than features. **This was further evidence that Martino's simulation considered the overlap of samples rather than features.** The code that was described at L123-125 was repeated as L135-138, using a standard deviation of 2 rather than 1. The resulting matrix, `B`, had 500 rows and 120 columns. Each row of `B` was identical with the columns being the value of the probability distribution function at 5.0 (i.e., `gradient` as defined at L131) for a normal distribution with the 120 values of `mu` as the mean and 2.0 (i.e., `sigma`) as the standard deviation. At L145, this matrix was used to replace the values in the upper left quadrant of `mat` in L139-145. The values of the first 500 rows in the last 80 columns were all nearly zero and were taken from the value of `mat` generated at L125.

The code shown below was similar to what was done at L131-138.

```
161         upper_col = int(upper_col - overlap)
162         # Make last block fill in the remainder
163         gradient = np.linspace(5, 5, nrows - upper_row)
164         mu = np.linspace(0, 10, ncols - upper_col)
165         sigma = 4
166         xs = [norm.pdf(gradient, loc=mu[i], scale=sigma)
```

```
167             for i in range(len(mu))]
168     B = np.vstack(xs).T * maxval
169 
170     mat[upper_row:, upper_col:] = B
```

The only difference was that in this code chunk they used a standard deviation of 4.0 (i.e., `sigma`) instead of 2.0. The new value of `B` defined at L168 was used to complete the value of `mat` by filling in the last 500 rows and the last 120 columns (L170). Again, the values of the first 80 columns in the last 500 rows were nearly zero.

In conclusion, there were three issues with the code in the `block_diagonal_gaus` function. First, the code at the beginning of the function (L117-125) was extraneous. Second, and more important, the features had the same relative abundance within each sample. Finally, and perhaps most important, samples were shared across features rather than features being shared across samples. As written in the original Jupyter notebook, the at the end of the `block_diagonal_gaus` function, values of `ncols` set to 200, `nrows` to 1000, `nblocks` to 2, `overlap` to 20, and `maxval` to 2500 resulted in a 1000 row by 200 column matrix. Below is a heatmap depiction of the raw values in `mat` with lighter colors are lower abundance while darker colors are higher abundance.

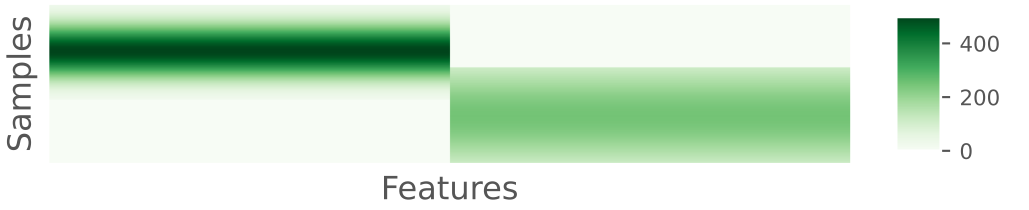

To highlight the observation that the probabilities were the same for each feature across non-overlapping samples within the same block, the following heatmap shows the relative abundances of each feature:

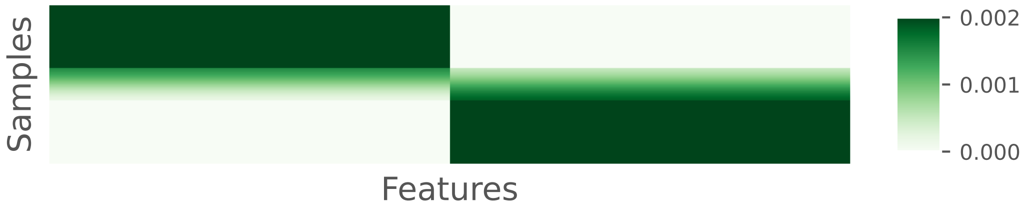

#### Review of `Homoscedastic`

The `Homoscedastic` function applied noise generated by randomly sampling from a normal distribution to each sample-taxon pair of the matrix using the abundance of the sample-taxon pair as the mean and `intensity` as the standard deviation. For the cases where overlap was included between the treatment groups, the intensity was the model parameter (e.g., 2500) divided by 15 (i.e., 166.6667) and for the case where there was no overlap between the treatment groups, the intensity was the model parameter divided by 60 (i.e., 41.6667).

```
42  def Homoscedastic(X_noise, intensity):
43      """ uniform normally dist. noise """
44      X_noise = np.array(X_noise)
45      err = intensity * np.ones_like(X_noise.copy())
46      X_noise = rand.normal(X_noise.copy(), err)
47
48      return X_noise
```

The code from L42-48 revealed another potential problem. The median abundance in `X_noise` coming into `Homoscedastic` across all samples was 115.7 with a maximum value of 498.6 (see the two previous heatmaps). Considering that random numbers were drawn from distributions with a standard deviation of 166.67 it was likely that negative values would result. In fact, for this example, 28.5% of the sample-taxon pairs were negative. The normal distribution likely should have been truncated by turning negative values to zeroes. Below is a heatmap depiction of `X_noise` after running the `Homoscedastic` function on L257 of the `build_block_model` function in `simulations.py`.

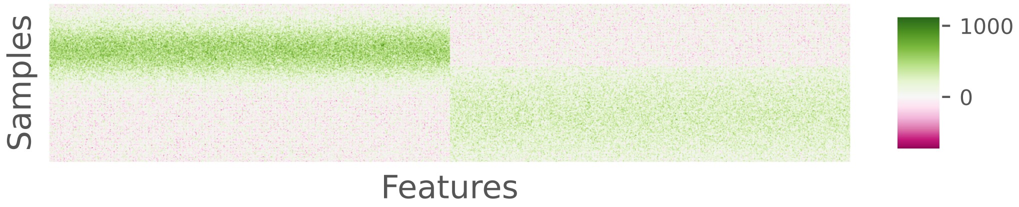

#### Review of `Heteroscedastic`

Similar to `Homoscedastic`, the `Heteroscedastic` function applied noise randomly sampled from a normal distribution using the abundance of the sample-taxon pair from `X_noise` as the mean and `intensity` as the standard deviation. Again, for the simulations of the partially overlapping features between treatment groups, the intensity was the model parameter (e.g., 2500) divided by 15 (i.e., 166.6667) and for the case where there was no overlap between the treatment groups, the intensity was the model parameter divided by 60 (i.e., 41.6667). The "Simulations" subsection of the "Materials and Methods" section of Martino indicated that the heteroscedastic source of noise was only supposed to be applied to a random subset of the sample-taxon pairs. However, review of the code for the `Heteroscedastic` function indicated that it applied noise to all values of `X_noise` rather than to a subset.

```
51  def Heteroscedastic(X_noise, intensity):
52      """ non-uniform normally dist. noise """
53      err = intensity * np.ones_like(X_noise)
54      i = rand.randint(0, err.shape[0], 5000)
55      j = rand.randint(0, err.shape[1], 5000)
56      err[i, j] = intensity
57      X_noise = abs(rand.normal(X_noise, err))
58
59      return X_noise
```

At L53 the code generated `err`, which was a matrix with the same dimensions as `X_noise` where every cell had the value of the intensity (e.g., 166.67). On L54-55 they selected 5000 random row and column indices that were then used in L56 to assign values of `err` to `intensity`. Except, L54-56 did not change the existing values of `err` since the values in the matrix were all already set to the value of `intensity` at L53. At L57, noise was added to all values of `X_noise` much like was done in `Homoscedastic` at L46. It was likely that the authors intended to use the numpy `zeros_like` function on L53 instead of the `ones_like` function. Then at L56, `err` would have had 5000 cells with the value of `intensity` and the `rand.normal` function would have changed only those values at L57. One notable difference to `Homoscedastic` was that L57 used `abs` to return the absolute value of `rand.normal`. Prior to applying the absolute value to the abundances with the additional noise, 32.2% of the sample-taxon pairs were negative. Using `abs` would certainly add noise to a subset of `X_noise`, but it would not be random since it would disproportionately impact low abundance features. As mentioned for `Homoscedastic`, returning zero rather than negative values or their absolute value would likely have been preferred. Below is a heatmap depiction of `X_noise` after running the `Heteroscedastic` function on L259 of the `build_block_model` function in `simulations.py`.

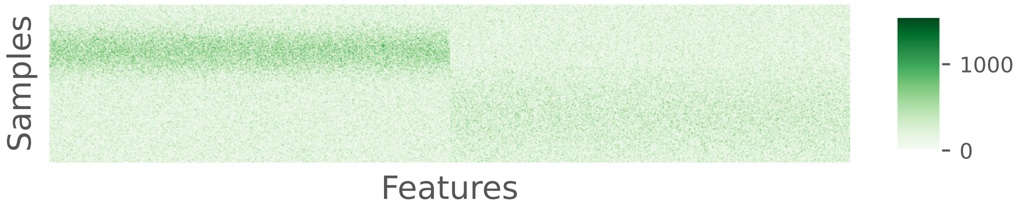

#### Review of `Subsample`

In the final step of `simulations.py`, the values of `X_noise` were used to generate a count matrix with the desired number of individuals per sample (L261) using the `Subsample` function.

```
51  def Subsample(X_noise, spar, num_samples):
52      """ yij ~ PLN( lambda_{ij}, /phi ) """
53      # subsample
54      mu = spar * closure(X_noise.T).T
55      X_noise = np.vstack([poisson(lognormal(np.log(mu[:, i]), 1))
56                           for i in range(num_samples)]).T
57      # add sparsity
58
59      return X_noise
```

This function took `X_noise` as input along with the `spar` and `num_samples` parameters. The value of `spar` was the same model parameter value that was used to scale the relative abundances in `block_diagonal_gaus` (e.g., 2500). At L54, the relative abundance of features within each sample was determined and then multiplied by `spar` to generate `mu` which was a `num_features` by `num_samples` array. These values were used to generate counts for each sample-taxon pair using a Poisson-lognormal distribution, which produced a new version of `X_noise`, which was returned to `build_block_model` and used for the rest of the analysis.

Below is a heatmap depiction of the central log ratio transformation of `X_noise` after adding 1 to each cell at the end of the `build_block_model` function in `simulations.py`.

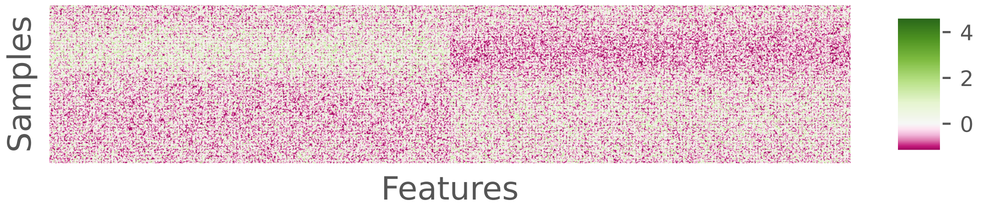

The matrix generated from `Subsample` did not have a consistent total number of counts for each sample. When using the model parameter of 2500, values near 4000 individuals per sample were obtained, but there was considerable variation. Using the model parameter value of 2500, the mean number of individuals per sample was 4131.765 with a range between 3421 and 4816. Considering the claim that RPCA is invariant to differences in sampling effort, it was desirable to control whether samples had the same number of individuals. As a workaround for this problem, I increased the model parameter (e.g., from 2500 to 2800) so that the minimum sample size was greater than the desired number of individuals and then down sampled each sample to the desired number.

##### Summary

Seven problems were found in the code as written in the original `simulations.py` script. They are listed here in order of importance.

**1. Samples, not features, were shared between the two treatment groups.** The intention was that features should be shared between treatment groups, not samples.
**2. Among samples from the same treatment group, the abundance distributions for each feature were uniform.** The result was that the relative abundance for all features in the same sample and treatment group had the same expected relative abundance. The intention was likely that the abundance distribution within each sample varied by feature.
**3. The total number of individuals varied across each sample.** Although there were claims that RPCA was insensitive to uneven sampling effort, this was not actually demonstrated in Martino. It was necessary for me to make the sampling effort uniform so this could be tested.
**4. The heteroscedastic source of noise was actually applied to all sample-taxon pairs.** The result was a second addition of noise to all sample-taxon values rather than the intended subset of values.
**5. Negative mean values were used as the mean in normal distribution sampling functions when calculating the heteroscedastic noise.** It was likely that the intention was for negative values to be transformed to zeros before sampling the distribution.
**6. The signs of negative values generated after applying the heteroscedastic noise were made positive.** It was likely that the intention was for negative values to be transformed to zeros.
**7. Setting the random number generator seed was inadequate to ensure that rerunning `simulations.py` with the same parameters yielded the same result.**

Taken together, these problems did not entirely invalidate the claims made by Martino about the performance of RPCA. Regardless, it was necessary to re-evaluate the claims with corrected code. Therefore, I rewrote `simulations.py` as an R script `simulations.R` (found in `workflow/scripts/`) that resolved the above-mentioned problems.

#### Review of simulation analysis code

Continuing my review of the code used in the published benchmarking analysis of the simulated data, I inspected the code that was used to calculate the PERMANOVA F statistics and P-values and k-Nearest

Neighbor (KNN) accuracy values. Review of the code in `deicode_benchmarking/simulations/negative_control.ipynb` and `deicode_benchmarking/simulations/scripts/cluster_build.py` and the data outputted to `deicode_benchmarking/simulations/cluster_models/simulation_subsampled_noisy.csv.gz` made it clear that each simulation (i.e., negative, positive, and overlapping) was only replicated a single time. This would have made it impossible to assess the sensitivity of the simulation results on the choice of the random number generator seed.

Aside from the lack of simulation replication, my primary critique concerned the KNN analysis. The manuscript stated “ordination output was compared by permutational multivariate analysis of variance (PERMANOVA) F-statistic and supervised k-nearest neighbor (KNN) classification cross-validation (40:60) split”. The convention for describing cross-validation splits is to indicate the percentage of samples used in the training set followed by the percentage used in the testing set. Therefore, according to a 40:60 split, 40% of the samples would have been used to train the model and 60% would have been used to test the model. Such a split would have been an odd choice since the training set is typically larger than the testing set. Furthermore, a split is often performed multiple times to remove bias from the selection of the random number generator seed.

Analysis of the benchmarking code made it clear that a 40:60 split was not used. In fact, for the negative and positive control analyses reported in their Table S1, the authors used 80:20 training-testing splits. This can be seen on L7 of box 186 from `deicode_benchmarking/simulations/negative_control.ipynb` shown here:

```
3   pcoa_tmp = pcoa(DistanceMatrix(distance.cdist(U_,U_))).samples
4   pcoa_tmp.index = T_.T.index
5   # split
6   X_train, X_test, y_train, y_test = train_test_split(pcoa_tmp, meta['group'].ravel(),
7                                                       test_size=.2,
8                                                       stratify=meta['group'].ravel(),
9                                                       random_state=42)
10  knn = KNeighborsClassifier(n_neighbors=10).fit(X_train, y_train)
```

In contrast, to analyze the data simulated with overlapping features they used 60:40 training-testing splits. This can be seen on L75 of `deicode_benchmarking/simulations/scripts/cluster_results.py` as shown here:

```
71  pcoa_tmp=pcoa(DistanceMatrix(distance.cdist(U_tmp,U_tmp))).samples
72  pcoa_tmp.index=subtmp_sub.index
73  # split
74  X_train, X_test, y_train, y_test = train_test_split(pcoa_tmp, meta['group'].ravel(),
```

```
75                                                        test_size=.4,
76                                                        stratify=meta['group'].ravel(),
77                                                        random_state=42)
78  knn = KNeighborsClassifier(n_neighbors=13).fit(X_train, y_train)
```

Review of the code in both `negative_control.ipynb` and `cluster_results.py` made it clear that only a single split of the data was performed.

The choice of the number of neighbors (i.e., hyperparameter k) used in the KNN analysis was not described in the manuscript. Often the training set will be repeatedly split to fit such hyperparameters (2). As shown above in the code at L10 from box 186 from `negative_control.ipynb`, a k value of 10 was used to classify the positive and negative control data. In contrast, as shown above in the code from `cluster_results.py` at L78, a k value of 13 was used to classify the samples with overlapping features.

In summary, the authors used inconsistent splits and k-values in their simulations, misstated the nature of the split, did not incorporate multiple splits, and provided no information for how the k-value was selected. These problems raised the question of whether their models were overfit (i.e., too small of a k value) or under fit (i.e., too large of a k value). Therefore, I implemented my own version of a KNN classifier (`workflow/scripts/test_train_knn.R`) that allowed me to input a distance matrix, training, validation, and testing fractions, and the number of iterations.

### Review of case study analysis code

#### Processing of case study datasets

After benchmarking RPCA using simulated data, Martino evaluated the method relative to using Bray-Curtis and weighted UniFrac distances with the Sponges and Sleep Apnea datasets (3, 4). Both datasets consisted of two treatment groups represented by varying numbers of samples, each with a wide distribution in the number of sequences (**Figure 5**). The datasets were analyzed separately by similar methods. Before commenting on the analysis of these datasets, several issues are noteworthy.

First, to create balanced experimental designs, the authors randomly selected an equal number of samples from each treatment group. While unbalanced design is problematic for KNN classification, PERMANOVA is not known to be sensitive to such designs. To mitigate the effects of the random number generator seed, they repeated this sampling. The text of the manuscript states that they performed “10-fold random subsamples of the data". In fact, at L45 of `deicode_benchmarking/case_studies/scripts/subsample.py` it was clear that they only performed 9-fold random subsamples.

```
44 # repeat randomized sub-sample 10 fold times
45 for fold_ in range(1,10):
46     # Randomly Sub-sample to balance groups
```

The python `range(start, stop)` function outputs values from the value of `start` and ends at one less than `stop`. For example:

```
1 list(range(1, 10))
```

```
[1, 2, 3, 4, 5, 6, 7, 8, 9]
```

Admittedly, this was unlikely to have had a meaningful impact on the results of the case study analysis.

Second, Bray-Curtis and weighted UniFrac distances are known to be sensitive to uneven sampling effort of the samples being compared; Martino claimed that RPCA was not. To mitigate the effects of uneven sequencing depth among samples, Martino performed a single subsampling of the data down to 1,000 sequences per sample when calculating the Bray-Curtis and weighted UniFrac distances; all samples with more than 1,000 sequences were used for RPCA without subsampling to a common number of sequences. This was encoded at L38-49 and L61-62 from `deicode_benchmarking/case_studies/scripts/distances.py`.

```
33 table_path = os.path.join(dir_path,subpath_,sub_set,'table.biom')
34 metadata_path_qiime = os.path.join(dir_path,subpath_,sub_set,'metadata.tsv')
35 # make qiime2 table form biom
36 table_path_qiime=os.path.join(dir_path,subpath_,sub_set,'table.qza')
37 subprocess.call("qiime tools import --input-path %s --type 'FeatureTable[Frequency]' --source-format BIOMV210Format --output-path %s"%(table_path,table_path_qiime), shell=True)
38 # sequencing depth cleaning (1000 read/sample)
39 subprocess.call("qiime feature-table filter-samples --i-table %s --p-min-frequency 1000 --o-filtered-table %s"%(table_path_qiime,table_path_qiime), shell=True)
40 # trim biom of any tree issues (should not be any but why not)
41 subprocess.call("qiime phylogeny filter-table --i-table %s --i-tree %s --o-filtered-table %s"%(table_path_qiime,tree_path_qiime,table_path_qiime), shell=True)
42 # rarefy (1,000)
43 subprocess.call("qiime feature-table rarefy --i-table %s --p-sampling-depth 1000 --o-rarefied-table %s"%(table_path_qiime,table_path_qiime), shell=True)
44 # bray-curtis
45 bray_dist = os.path.join(dir_path,subpath_,sub_set,'Bray_Distance.qza')
46 subprocess.call("qiime diversity beta --i-table %s --p-metric 'braycurtis' --o-distance-matrix %s"%(table_path_qiime,bray_dist), shell=True)
47 # generalized weighted alpha=1.0
48 wone_dist = os.path.join(dir_path,subpath_,sub_set,'GUniFrac_alpha_one_Distance.qza')
```

```
49 subprocess.call("qiime diversity beta-phylogenetic-alt --i-table %s --i-phylogeny %s --p-metric generalized_unifrac --p-alpha 1.0 --o-distance-matrix %s"%(table_path_qiime,tree_path_qiime,wone_dist), shell=True)
```

```
61 # run RPCA
62 subprocess.call("deicode_rpca --in_biom %s --output_dir %s --rank %s --min_sample_depth 1000"%(table_path,sub_path_sub,str(ranks[dataset_])), shell=True)
```

From this code, one will note on that `table_path_qiime` was created on L37 and updated at L39 (filtering to remove samples smaller than 1,000 sequences), L41 (tree cleaning), and L43 (each sample subsampled once to 1,000 sequences). The data in `table_path_qiime` was used to calculate Bray-Curtis and UniFrac distances at L46 and L49, respectively. When RPCA was performed on L62, the input was `table_path`, which was created on L33 and used as the input to create the biom-formatted file in the initial version of the data in `table_path_qiime` (L37). It is also worth noting that Bray-Curtis and weighted UniFrac distances from a single subsampling were used rather than using the average of a large number of subsamplings (e.g., 100) (5).

As shown in **Figure 5**, 1,000 appears to have been an arbitrary threshold. It could have been greater than 10,000 sequences for both datasets without losing a significant number of samples. That so many more sequences were included in the RPCA analyses could create an advantage to the sensitivity of RPCA over the other distance calculations.

Third, in my attempts to reproduce the weighted UniFrac distances, I found that the trees provided in `deicode_benchmarking/case_studies/data/*/tree_relabelled.tre.gz` did not include all of the sequences included in the count data found in the author-supplied biom files. In the Sleep Apnea dataset the biom file contained 1369 OTUs and the Sponges biom file had $1.116 \times 10^4$ OTUs. The tree files contained 1345 and $1.1074 \times 10^4$ OTUs, respectively. When accounting for the total number of sequences represented by each OTU, 39.7115899 of all of the sequences were missing from the Sleep Apnea trees and 0.6888975 were missing from the Sponges trees. Martino removed sequences that were missing from the OTU table at L39. The comment preceding this line, “trim biom of any tree issues (should not be any but why not)” seems to indicate that they did not anticipate any discrepencies between the two types of data. When I removed this line from the code, I received an error message that stated, “Compute failed in one_off: Table observation IDs are not a subset of the tree tips.”. Given the large percentage of total reads removed from the Sleep Apnea dataset, the discrepency would have the greatest effect on the Bray-Curtis distances from that dataset. Since the filtered OTU table was not used as input to RPCA, this resulted in another uneven comparison between the methods.

Fourth, the methods section of Martino stated, “Both data sets were then preprocessed with the robust centered log ratio (rclr) transform, and RPCA was run with a rank of 2 because there were two metadata categories of interest in each comparison”. However, different ranks were used when running RPCA for the Sleep Apnea and Sponges dataset. Three ranks were used with the Sleep Apnea dataset and two were used with the Sponges dataset. This can be seen at L23 and L24 of `deicode_benchmarking/case_studies/scripts/distances.py`

```
23 ranks['Sleep_Apnea'] = 3 # three clusters here
24 ranks['Sponges'] = 2 # two clusters here
```

As indicated above at L62 of the same file, the value of `ranks` set here in L23 and L24 were given to `deicode_rpca` in the `--rank` argument for each dataset. That a rank of 3 was used for the Sleep Apnea dataset was used can also be seen in Martino’s Figure 4F. The percent of the total variation explained by the two axes did not add to 100% for the RPCA data. In contrast, for the Sponges dataset shown in Martino’s Figure 4E they add to 100%.

Among these issues, the most salient issue was likely the disparity in the number of sequence reads used by RPCA and in calculating the Bray-Curtis and weighted UniFrac distances.

#### PERMANOVA analysis of case study datasets

The factors outlined in the previous section would likely also impact the results of Martino’s PERMANOVA analysis of the case study datasets. Several additional factors were worth raising and emphasizing.

First, throughout Martino’s analysis, they reported the PERMANOVA F-statistic rather than the test’s P-value. This was likely because there was a large effect size between the treatment groups in both studies that was clearly evident in the first two axes of the ordinations shown in Martino’s Figure 4EF. Reporting all tests as having significant P-values regardless of the method used to calculate the distance would not have been interesting. However, comparing F-statistics was biased in favor of RPCA. This was because RPCA was run using 2 (i.e., Sponges) or 3 (i.e., Sleep Apnea) axes and then used as the basis for calculating the Euclidean distance between samples. The analysis effectively removed higher ordered components thus dramatically reducing the $MS_{error}$ term in the denominator of the F-statistic calculation. In contrast, Bray-Curtis and weighted UniFrac distances were used without further processing in PERMANOVA leaving in the higher ordered components. As done in my analysis, using datasets where the first two axes for Bray-Curtis distances did not explain so much of the variation would have provided a more balanced comparison of the methods.

Second, Martino's Figure 4A-E did not show the variation across their 9 subsamplings for their PERMANOVA or KNN classification analyses. This can be seen in L43-72 of code block 104 from `deicode_benchmarking/case_studies/case_studies.ipynb`. An illustrative example is shown here:

```
43  # plot benchmarking
44  fstat_ax1.set_title('PERMANOVA F-statistic', fontsize=fontsize_)
45  sns.pointplot(x='N-Samples',y='Values',hue='Method',
46                data=both_perm_res['Sponges'].sort_values('Method',ascending=False),
47                palette=colors_, ci=0, ax=fstat_ax1)
```

The `pointplot()` function from the Seaborn data visualization library plots the mean of multiple values across a series of categorical values. By default it would have returned the 95% confidence interval. However, the authors removed the confidence intervals when they used the `ci=0` argument in L47, which suppressed its output. Using the data included in the `deicode_benchmarking` repository `deicode_benchmarking/case_studies/subsample_results.tar.gz`, I calculated the median, interquartile range (IQR), minimum, and maximum values across the 9 subsamplings for each dataset, distance method, and type of analysis. The conditions where 79 samples were taken from each treatment in the Sponges dataset and 92 samples were taken from each sample in the Sleep Apnea dataset were used in this analysis.

| analysis | dataset | name | median | iqr | min | max |
|---|---|---|---|---|---|---|
| KNN | Sleep_Apnea | Bray_Curtis | 0.82 | 0.03 | 0.80 | 0.84 |
| KNN | Sleep_Apnea | GUniFrac_Alpha_One | 0.43 | 0.01 | 0.42 | 0.46 |
| KNN | Sleep_Apnea | Robust_Aitchison | 0.86 | 0.00 | 0.86 | 0.86 |
| KNN | Sponges | Bray_Curtis | 0.85 | 0.02 | 0.80 | 0.90 |
| KNN | Sponges | GUniFrac_Alpha_One | 0.84 | 0.03 | 0.80 | 0.89 |
| KNN | Sponges | Robust_Aitchison | 0.91 | 0.02 | 0.89 | 0.94 |
| PERMANOVA | Sleep_Apnea | Bray_Curtis | 18.15 | 0.28 | 17.75 | 18.42 |
| PERMANOVA | Sleep_Apnea | GUniFrac_Alpha_One | 9.51 | 0.15 | 9.25 | 9.80 |
| PERMANOVA | Sleep_Apnea | Robust_Aitchison | 79.78 | 0.00 | 79.78 | 79.78 |
| PERMANOVA | Sponges | Bray_Curtis | 72.82 | 8.27 | 67.13 | 78.33 |
| PERMANOVA | Sponges | GUniFrac_Alpha_One | 133.47 | 11.24 | 121.50 | 150.84 |
| PERMANOVA | Sponges | Robust_Aitchison | 353.42 | 70.73 | 313.17 | 472.70 |

These results indicate a modest amount of variation between the replicates. The RPCA data for the Sleep Apnea dataset had no variation because all samples were used in the analysis and no subsampling was performed to obtain equal sampling depth across the samples.

Third, the performance of PERMANOVA is not known to be negatively impacted by unbalanced designs (i.e., where treatment groups have different numbers of samples). Although understanding the effect of the number of samples on PERMANOVA's statistical power is a worthy question, it was curious that the entire Sponges dataset was not included in Martino's analysis. The Sponges dataset had 124 Healthy samples and 79 Stressed samples with more than 1,000 sequences. Yet, for the largest number of samples in Figure 4 the largest subsample was 79 Healthy and 79 Stressed samples.

Among the issues I have identified, the primary concerns with how Martino used PERMANOVA with the case studies were (i) that they used datasets with large effect sizes and (ii) that RPCA has a built-in bias over Bray-Curtis and weighted UniFrac distances for such datasets when calculating the F-statistics, which were all significant regardless of the magnitude of the statistic.

#### KNN analysis of case study datasets

In addition to PERMANOVA, Martino also used KNN classification to demonstrate the performance of RPCA over weighted UniFrac and Bray-Curtis distances with their two case study datasets. The issues identified earlier with the simulation data were also present with their case study analysis (i.e., 60:40 vs. 40:60 splits and choice of K). Several additional issues were identified in the application of KNN to the case study datasets. The Python code that was used to implement their KNN analysis was found in `deicode_benchmarking/case_studies/scripts/sub_results.py` between L123 and L174.

```
123 for dataset_,subs in distances.items():
124   nn_res[dataset_]={}
125   nn_res_tmp[dataset_]={}
126   # set splits for each dataset here
127   max_size=np.max([j for i,j in distances[dataset_].keys()])
128   dist_tmp=distances[dataset_][(1,max_size)]['Bray_Curtis'] #any method to split
129   meta_=meta[dataset_][(1,max_size)]['metadata']
130   pcoa_tmp=pcoa(DistanceMatrix(dist_tmp)).samples
131   pcoa_tmp.index=dist_tmp.index
132   # prepare binary classifier
133   y=meta_[case_study[dataset_]['factor']].values
134   proc=preprocessing.LabelBinarizer()
135   proc.fit(y)
136   meta_['y_encode']=proc.transform(y)
137   # split
138   X_train, X_test, y_train, y_test = train_test_split(pcoa_tmp, meta_['y_encode'].ravel(),
139                                                       test_size=.4,
140                                                       stratify=meta_['y_encode'].ravel(),
141                                                       random_state=42)
142   train_index =X_train.index
143   test_index = X_test.index
144   # run KNN for every subset with train and test splits
145   for (fold_,Nsamp_),methods_ in subs.items():
146     meta_=meta[dataset_][(fold_,Nsamp_)]['metadata']
147     if len(meta_.index)<Nsamp_:
148         continue
149     nn_res[dataset_][(fold_,Nsamp_)]={}
150     nn_res_tmp[dataset_][(fold_,Nsamp_)]={}
```

```
151      for count_,(method,dist_tmp) in enumerate(methods_.items()):
152        if method in ['Bray_Curtis','GUniFrac_Alpha_One', 'Robust_Aitchison']:
153          nn_res[dataset_][(fold_,Nsamp_)][method]={}
154          meta_=meta[dataset_][(fold_,Nsamp_)]['metadata']
155          tmp_train_index=list(set(train_index)&set(meta_.index))
156          tmp_test_index=list(set(test_index)&set(meta_.index))
157          # encode
158          meta_['y_encode']=proc.transform(meta_[case_study[dataset_]['factor']].values).ravel()
159          pcoa_tmp=pcoa(DistanceMatrix(dist_tmp)).samples[['PC1','PC2']]
160          pcoa_tmp.index=meta_.index
161          # Run default classifier
162          acc_s=[]
163          for fold_rep in range(11):
164            k_=int(min(meta_.loc[tmp_test_index,:].shape)/1.4)
165            knn = KNeighborsClassifier(n_neighbors=k_).fit(pcoa_tmp.loc[tmp_train_index,:],
166
    ↪  meta_.loc[tmp_train_index,:]['y_encode'].astype(int).ravel())
167            acc_ = accuracy_score(meta_.loc[tmp_test_index,:]['y_encode'].ravel(),
168                                  knn.predict(pcoa_tmp.loc[tmp_test_index,:]).astype(int))
169            acc_s.append(acc_)
170
171          nn_res[dataset_][(fold_,Nsamp_)][method]['R^{2}'] = np.mean(acc_)
172          nn_res_tmp[dataset_][(fold_,Nsamp_)]=pd.DataFrame(nn_res[dataset_][(fold_,Nsamp_)])
173
174    both_nn[dataset_]=pd.concat(nn_res_tmp[dataset_])
```

First, when describing their KNN analysis Martino again stated that they performed 40:60 splits. As shown at L139 the test fraction was 0.4 rather than 0.6 as would have been expected from a 40:60 split. Martino actually implemented a 60:40 split where 60% of the data were used to train the model an 40% were used to test the model.

Second, inspection of the above code chunk demonstrated that they only performed a single split of the data at L138-141. Moreover, there was only one split across all subsamplings for each dataset. At L142 and L143 they assigned the output of the training-testing split from L138-141 to two variables, `train_index` (L142) and `test_index` (L143). These variables contained the indices for the samples in the training and testing set across all the samples, stratified by the number of samples in each treatment group (L140). It is important to note that the code between L124 and L143 was only run once for each dataset creating a single 60:40 split that was then used for each subsampling. On L155 and L156, training and testing sets, respectively, were created for each subsample based on the split that was performed at L138-L143. Because the sample subsampling was not performed based on the 60:40 split, different numbers of samples from each treatment group appeared in each of the replicate subsamples. Thus, the training and testing sets were not equally represented by both treatment groups. Considering KNN is sensitive to unbalanced sampling designs, this would have artificially increased or decreased the accuracy of the classification. Finally, at L163 to L169 it appeared that KNN was run 10 times (i.e., see the `range(11)` statement). Because the training and testing data were the same in each iteration of the loop, the values of `acc_` that were saved into `acc_s` at L169 were identical. For each dataset, subsampling, and iteration of subsampling Martino should have split the data into a training and testing split stratified by the treatment group. Instead they did the split once on the entire dataset. Furthermore, these splits should have been repeated multiple times (e.g., 10 or 100) with the mean accuracy of those repeats returned as the mean accuracy for the iteration of the subsampling for the dataset. The lack of replication or balanced stratification of the treatment groups within the subsamples significantly compromised the reliability of the KNN classification accuracy results.

Third, Martino did not fit the hyperparameter, k, to their data and the code to select a value for k was clearly incorrect. The value of k was calculated at L158

```
158 k_=int(min(meta_.loc[tmp_test_index,:].shape)/1.4)
```

Martino subsetted the metadata data frame (i.e., `meta_`) using the values in the test data (i.e., `tmp_test_index`). Then they used the `.shape` function to return the number of rows and columns. The smaller number was then divided by 1.4 and converted to an integer. There were several problems with this calculation. First, one would have expected the choice of k to be dependent on the number of samples included in the **training** dataset and not the **testing** dataset since the observations in the training and not the testing dataset affected the classification. Instead Martino subsetted `meta_` by the samples in the **testing** dataset. Furthermore, the number of samples and not the amount of metadata for each sample would have been relevant to fitting the value of k. To illustrate this further, the number of columns for the Sponges dataset was 46 and for the Sleep Apnea dataset it was 50 (one column added for `y_encode`). Therefore, when more than 115 and 125 total samples, respectively, were in a subsample k would have been selected based on the number of columns in `meta_` (e.g. 46/0.4 = 114). These would have been the 150 and 158 subsamples for the Sponges datset and 150 and 184 subsamples for the Sleep Apnea dataset. The values of k would have been 32 and 35 for these subsamples in the Sponges and Sleep Apnea datasets, respectively. For the other subsamples, k would have been the total number of samples in the subsample times 0.4 divided by 1.4. The subsamples with 70 samples would have had a k value of 20 and the 110 subsamples would have had a k value of 31. Because the stratification was not balanced (see above discussion) these thresholds were approximate. For context, in the simulations there were 200 samples with 120 in the training set and 80 in the testing set. A k of 13 was used. Therefore, the ratio of k to the number of samples in the training set was 0.108333. Although no justification was given for the k of 13, if the same ratio had been used with the case study datasets, the k values would have been 4, 7, 9, 10, and 12 for the subsamples of 70, 110, 150, 158, 185 samples, respectively (e.g., 150 times 0.6 times 0.108333 = 9.75, converted to an integer gives 9). As discussed above with the KNN analysis of the simulation data, a preferred approach would have subdivided the training dataset into a secondary training set and a validation datset. The secondary and validation training sets could have been used to identify the optimal value of k to use with the full training set on the testing dataset (2). As described above with the discussion of the choice of k for the simulation data, if a too small value of k were chosen the model would have been overfit to the data while too large of a value of k could have underfit the model to the data. Given the problems identified here, it was clear that there was no clear logic to the choice of k. Therefore, it was difficult to have confidence in the reported accuracy values from the KNN analysis.

Fourth, at L159 two dimensions from a principle coordinates analysis (PCoA) performed on the Bray-Curtis and weighted UniFrac distances and two dimensions from the RPCA were used to define the dataset for KNN; this line would have ignored the third dimension from the Sleep Apnea dataset and many dimensions from the PCoA. The KNN used a Euclidean distance to find the k nearest neighbors in the training dataset for each value in the testing dataset. Because so much information in the higher ordination axes was censored from the Bray-Curtis and weighted UniFrac distances, those methods were disadvantaged relative to RPCA. Considering `KNeighborsClassifier` could take distance matrices as input, it was surprising that Martino did not use the Bray-Curtis and weighted UniFrac distances to make a more fair comparison to RPCA. Overall, the implementation of the KNN analysis was biased towards favoring the RPCA approach. This was exacerbated by the choice of datasets that had large differences between their treatment groups.

Given the problems in the implementation of KNN by Martino, I rewrote the method in R. The functions to run my analysis were found in `workflow/scripts/test_train_knn.R`. I rewrote the KNN analysis to implement the fitting of k using the training dataset with a secondary training and validation dataset. I was able to alter the training, testing, and validation fractions and number of replications. I could use my R code to replicate Martino's approach or the approach outlined by Topcuoglu et al. (2).
